# PFTK1 kinase regulates axogenesis during development via RhoA activation

**DOI:** 10.1101/2022.01.11.475789

**Authors:** Yasmilde Rodríguez González, Fatemeh Kamkar, Paymaan Jafar-nejad, Suzi Wang, Dianbo Qu, Leticia Sanchez Alvarez, Dina Hawari, Margaret Sonnenfeld, Ruth S. Slack, Paul Albert, David S. Park, Alvin Joselin

## Abstract

PFTK1/Eip63E is a member of the Cyclin-dependent kinases (CDKs) family and plays an important role in normal cell cycle progression. Eip63E expresses primarily in postnatal and adult nervous system in Drosophila melanogaster but its role in CNS development remains unknown. We sought to understand the function of Eip63E in the CNS by studying the fly ventral nerve cord during development. Our results demonstrate that Eip63E regulates axogenesis in neurons and its deficiency leads to neuronal defects. Functional interaction studies performed using the same system identify an interaction between Eip63E and the small GTPase Rho1. Furthermore, deficiency of Eip63E homolog in mice, PFTK1, in a newly generated PFTK1 knockout mice results in increased axonal outgrowth confirming that the developmental defects observed in the fly model are due to defects in axogenesis. Importantly, RhoA phosphorylation and activity is affected by PFTK1 in primary neuronal cultures. We here report that GDP bound inactive RhoA is a substrate of PFTK1 and PFTK1 phosphorylation is required for RhoA activity. In conclusion, our work establishes an unreported neuronal role of PFTK1 in axon development mediated by phosphorylation and activation of GDP-bound RhoA. The results presented add to our understanding of the role of Cdks in the maintenance of RhoA mediated axon growth and its impact on CNS development and axonal regeneration.

## Introduction

Cyclin Dependent Kinase (CDK) are involved in the regulation of several cellular processes including cell cycle regulation, transcription, translation and neuronal differentiation and function (Malumbres and Barbacid, 2005). Several of these kinases and their interaction partners are typically highly expressed and active in neuronal cells. This includes CDK5, CDK16-18 (PCTAIRE 1-3/ PCTK1-3) and CDK14-15 (PFTAIRE/ PFTK1-2). The best characterized of these CDKs, CDK5, is predominantly active in neurons owing to the tissue-specific expression of its activators, p35/25 and p39. CDK5 deficiency in mice leads to perinatal lethality as a result of defective neuronal migration, differentiation and survival. Both PFTK1 and PCTK1 have been shown to be activated by cyclin Y (Jiang et al., 2009; Mikolcevic P, 2012). PFTK1, first identified in *Drosophila melanogaster* (Dmel) (Sauer et al., 1996), and later in mice (Lazzaro and Julien, 1997) and humans (Yang and Chen, 2001), is highly expressed in the Central Nervous System (CNS) (Besset et al., 1998; Lazzaro et al., 1997), predominantly in the brain and likely plays an important role in development since PFTAIRE deficient flies die mostly at early larval stages (Stowers et al., 2000). Notwithstanding, the function of PFTK1 in CNS development and the downstream substrates PFTK1 regulates has remained largely undiscovered.

In mammals, PFTK exists as 2 separate genes: *PFTAIRE1/PFTK1/CDK14* (Yang and Chen, 2001) and *PFTAIRE2/PFTK2/Als2cr7/CDK15* (PubMed Gene ID: 65061). PFTK1 is highly conserved among different species from yeast to mammals (Liu and Kipreos, 2000). The fly and worm genome code for the gene known as Eip63-Ecdysone-induced protein 63E which shares 70% homology to mouse PFTK1 (Liu and Kipreos, 2000; Stowers *et al*., 2000). The human *PFTK1* and mouse *Pftk1* are located at (7q21-q22) and (5 A1; 2.61 cM) respectively (Lazzaro and Julien, 1997) and show 96% homology. *PFTAIRE2/ PFTK2/ Als2cr7/ CDK15* is recognized in GenBank database, and has been reported for its role in conferring resistance to tumor necrosis factor-related apoptosis-inducing ligand (TRAIL) in cancer cells (Park et al., 2014). In the adult mouse, *Pftk1* mRNA is located both in neuroglia and neurons (Besset *et al*., 1998; Lazzaro *et al*., 1997) in different regions of the brain such as cortex, hippocampus, thalamus, cerebellum, and in the spine. PFTK1 is localized in neuronal cell bodies but not in neurite extensions. During mouse development, *Pftk1* mRNA is expressed at embryonic days 12.5, 15.5, and even 18.5 but soon rises after birth, at P 10.5, and reaches a maximum, comparable to levels in adult mice. This is different from the expression pattern of CDK5 that has a continuous increase from embryonic day 12, followed by a significant increase at birth (Lazzaro *et al*., 1997). All this evidence, together with the knowledge that PFTK1 and CDK5 share a high amino acid sequence similarity (∼ 50-52%), led us to examine the role of PFTK1 during CNS development.

We first examined the role of PFTK1 homolog Eip63E in Dmel ventral nerve cord (VNC) in the regulation of axon and neuronal structure. We show that Eip63E regulates axogenesis in neurons and Eip63E deficiency or downregulation leads to defects in the axon and neuronal structure of the VNC. Furthermore, our functional interaction studies using this model led us to identify Rho1 as important mediator of Eip63E function. To further confirm this role of PFTK1 in CNS, we describe the characterization of a *Pftk1* knockout (KO) mice. Although, no gross abnormalities were observed in morphology, fertility, life span or anatomical brain structures of the PFTK1 KO mice within the parameters tested, perturbations in axonal growth were observed in cultured cortical neurons. Corroborating our observations in the flies, overexpression of a dominant negative PFTK1 in WT cultured cortical neurons and primary cortical neurons from *Pftk1* KO mice, exhibit faster-growing axons. In line with our observations in the flies, investigations show compelling evidence that PFTK1 phosphorylates GDP-RhoA, and this leads to RhoA activation. These data together point to an important and unexplored role of PFTK1 in axogenesis through its interaction with RhoA implicating PFTK1s role in CNS development.

## Results

### Eip63E KO embryos show defasciculation and misguidance of axons in VNC

To examine the role of PFTK1 in CNS development, we used the VNC of Dmel embryos, since it provides an excellent model to study developmental processes (Garbe and Bashaw, 2004; Sanchez-Soriano et al., 2007). Additionally, flies only have one homolog gene for PFTK (Eip63E) and do not have other neuronal CDKs closely related to PFTK1 that could add a level of complexity, due to possible compensation.

We used two different Eip63E mutant Dmel lines, carrying the Df(3L)E1 and Eip63E^81^ alleles respectively (Stowers *et al*., 2000). The nature of both these alleles is different in that the former is a result of a large deletion that spans the transcription initiation region to the entire kinase domain of Eip63E and affect two other adjacent genes, namely SDE3/Armitage (Cook et al., 2004) and 1(3)63D (Stowers *et al*., 2000). Alternatively, Eip63E^81^ is a smaller 48 bp in-frame deletion that removes 16 codons within Eip63E conserved kinase domain (213-228). Since both alleles leads to embryonic lethality, the fly’s populations are maintained as heterozygous mutants with the use of balancer chromosomes. Because we wanted to examine embryos, both mutant lines were balanced with the TM3krGFP,Sb balancer chromosome which expresses GFP under the control of Krüppell promoter from embryonic stage 9-10 to adulthood (Casso et al., 2000). Once the expression of GFP kicks in (absent before stage 9), Eip63E deficient embryos are distinguishable from the rest of the population by the absence of GFP expression.

Initially, BP102 or fasciclin-II (1D4) were both used as different markers to assess the VNC structures of stage 14-15 embryos. Both are expressed in the axon bundles that define the commissural and longitudinal connectives of the fly VNC (Fambrough et al., 1996; Hummel et al., 1999). Both markers confirmed axonal defects in the Eip63E KO embryos for both mutant alleles (Fig 1). KO embryos (GFP negative) show a diffuse appearance of the axonal pathways (Fig. 1A and 1B) suggesting defasciculation of axon bundles. Additionally, axons appear not to follow the appropriate pathways, leading to malformation of the ladder-like normal axonal structure of the VNC, with disruption of both commissural and longitudinal pathways (Fig. 1A and 1B).

**Figure 1.**
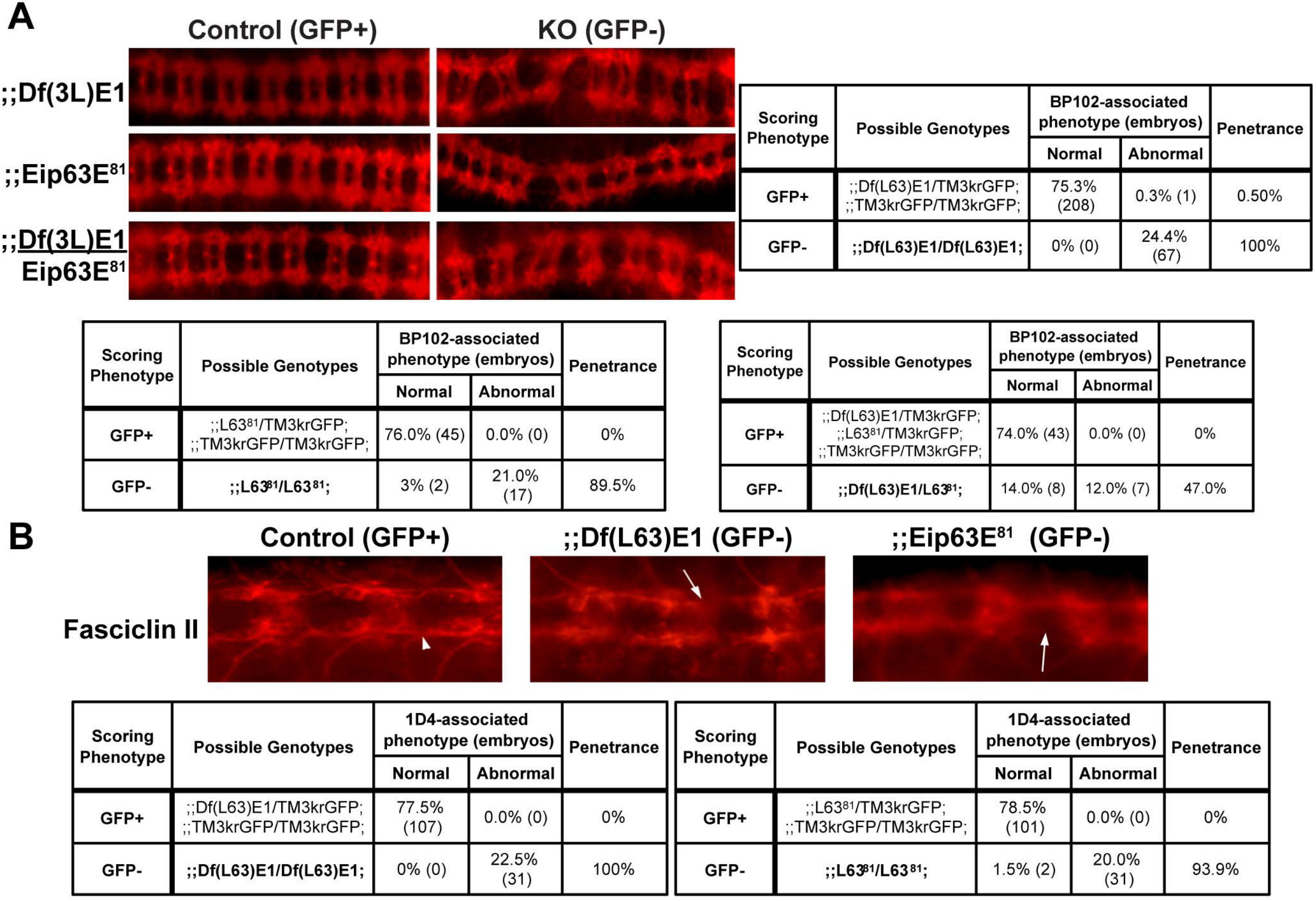
Eip63E deficiency leads to defasciculation and misguidance of VNC axons. Ventral views of whole mount stage 14-15 embryos from both Eip63E mutant lines, co-stained for GFP and BPI 02 (A) or Fasciclin-II (1D4 mAb) (B). Penetrance of axon abnormalities was calculated as the % of embryos of a given scoring phenotype that show any defect from the total population of such phenotype (GFP+ or GFP-embryos). Arrowhead points to normal looking longitudinal connectives and arrows to disrupted structures in KO embryos. L63^81^: Eip63E^81^.

Given the difference of genetic background between the mutant alleles, we needed to test if the observed VNC defects were primarily related to Eip63E. We therefore generated trans-heterozygous mutant embryos containing one copy of each of the mutant alleles and stained for BP102. As shown in Fig. 1A (bottom panel) and its corresponding scoring chart (Fig. 1A, bottom right), axonal defects similar to those of the single deficiencies were observed in the trans-heterozygous mutants. It should also be noted that no defects were ever observed in GFP-positive embryos, suggesting that none of the mutant alleles exhibit dominant-negative behavior. Together, these results suggest that axonal defects were indeed caused, at least in part, by Eip63E deficiency and not only due to background mutations.

### Early neurogenesis events are defective in Eip63E mutants

An early event in Dmel CNS development is the formation of the ventral midline. This has an essential impact in axon outgrowth/guidance and the proper ladder-like scaffold of axons of the VNC (Hummel *et al*., 1999; Klambt et al., 1991). To determine if midline defects were primary to the defects observed at stage 14-15, we assessed midline formation in embryos of the Eip63E^81^ mutant line as compared to wild type W^118^ embryos. W^118^ was needed as control in these studies since we did not have any way to genotype the Eip63E^81^ embryos before stage9 due to absence of GFP expression under the Kruppel promoter (balancer) before that stage (this also rules out GFP as a source of the defects). We performed immunostaining for single-minded (sim), a protein which is exclusively expressed in midline precursors and their progeny and is a master regulator of the differentiation of midline neurons and glial cells (Nambu et al., 1990). Eip63E deficiency does not have any apparent effect on midline formation, since no defects were found in embryos from the Eip63E^81^ line (Stages 5-10) compared to WT embryos (Fig. 2A). Furthermore, HRP staining at stage 11 (Fig. 2C, bottom panel) suggests that the ventral midline appears normal in Eip63E KO embryos even when the deficiency might lead to axon defects and abnormal HRP staining at later stages.

**Figure 2.**
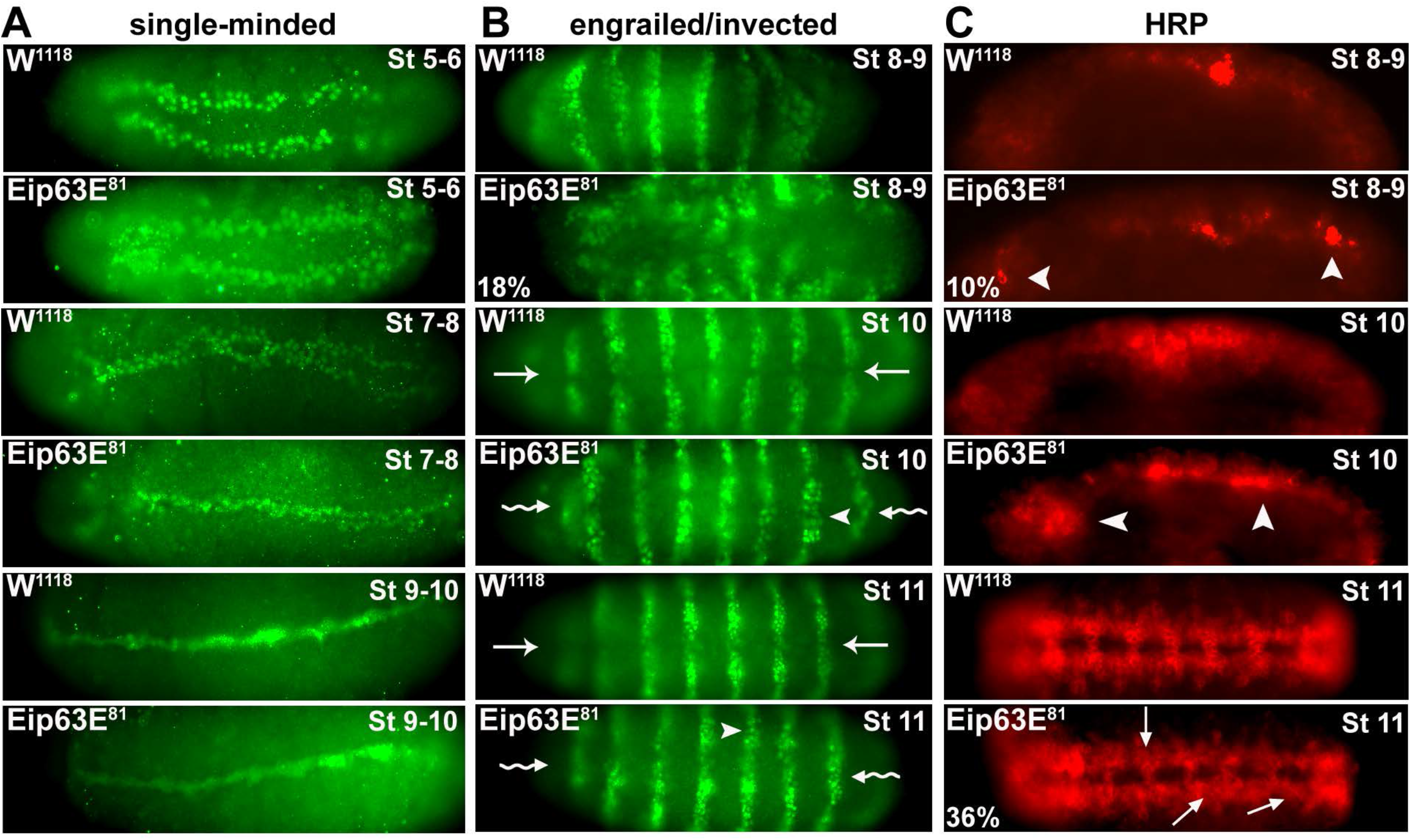
Eip63E deficiency leads to early embryonic developmental defects. Ventral views (unless otherwise specified) of whole mount embryos are shown head to the left. Stage 5-8 embryos are from Eip63E^81^/TM3krGFP,Sb line are of unknown genotype. Panels for stages 10-11 show Eip63E^81^ homozygous mutant embryos. (A) Staining for single-minded at early embryonic stages (st 5-10) show no gross difference in midline formation but cellular morphological differences are noticable. (B) Staining for engrailed/ invected (4D9 mAb). 18% (n=48) of total population of stage 8-9 embryos show acute segmentation defects. At stage 10-11, cell bodies appear disorganized (arrowheads) to the extent of loss of hemi-segments division at the midline (wavy arrow) compared to control (straight arrow). (C) Staining for the HRP antigen. In top 4 panels, a lateral view of embryos is shown. 10% (n=48) of st 8-9 embryos show premature HRP staining (arrowheads). At late stage 11, precocious axon outgrowth (arrows) seen in 36% of KO embryos (n=52).

Another hallmark of the Dmel embryo development is the segmentation of the embryo which is triggered early (within the first 3h) leading to the formation of stripe-like segments along the anterior-posterior axis. Segment polarity genes are part of the expression cascade that regulate this process and are expressed in 14 stripes at the onset of gastrulation (∼3h, stage 6-7) (Weigmann et al., 2003). These genes, which include engrailed (en) and invected (inv), are major players in neurogenesis (Bhat, 1999; Bhat and Schedl, 1997). We examined the expression pattern of en/inv proteins to evaluate if segmentation and/or neuroblasts delamination/segregation was affected in Eip63E KO embryos. Immunostaining with the 4D9 antibody, that recognizes both en/inv proteins, revealed that 18% of stage 8-9 embryos from the Eip63E^81^/TM3krGFP,Sb line show abnormal phenotypes (Fig. 2B). It is likely that embryos with affected segmentation (Fig. 2B, second panel down from the top) do not make it further into development. Interestingly, those that can compensate (stages 10-11) do not show segmentation defects but disorganization and morphological differences of cell bodies within segments. Since genotyping is not practical at these stages, it is not possible to link these phenotypes to Eip63E deficiency. It should be noted that comparable defects were observed in similarly timed embryos from the Df(3L)E1/TM3krGFP,Sb line (data not shown). In comparison, only 3% of WT embryos of similar stages show any defects. Although abnormalities were also observed in older KO embryos (stages 10), their incidence decreased (8% Eip63E^81^). This is likely due to the adverse effects of the defect on the survival of defective embryos combined with a decreased yield due to treatment methodology for the morphologically-defective embryos. Taken together, these results suggest a potential role for Eip63E in delamination of neuroblasts and/or their segregation.

### Axon outgrowth occurs prematurely in Eip63E KO embryos

Axogenesis in Dmel embryos starts at ∼7:30h into development (early stage 12) after specification of the midline cells that will guide the pioneer axons contra-laterally to cross the midline (Hummel *et al*., 1999; Jacobs and Goodman, 1989). To elucidate how early in CNS development axon defects start to appear, we examined CNS features using anti-horseradish peroxidase antibody which recognizes a carbohydrate epitope (called HRP for simplicity) present on several cell-surface molecules expressed by neuronal cell bodies and axons (Snow et al., 1987) including fasciclin II (Desai et al., 1994). 10% of stage 8-9 embryos mixed population from the Eip63E^81^ line (n=48) showed premature HRP staining distinct from WT W^118^ embryos (Fig. 2C, top 2 panels). Lateral views of the embryos are presented since this abnormality became more evident when looking laterally. Stage 9-10 W^118^ embryos show faint, background staining along the ventral region of the embryo where VNC axons would later appear. In contrast, Eip63E^81^ embryos show HRP-positive structures at stages 9-10 distinct from W^118^ embryos (Fig. 2C, middle 2 panels). By late stage 11, 36% of Eip63E^81^ KO embryos (GFP-, n=52) show premature, and in some cases misguided, axon outgrowth affecting several segments of the VNC (Fig. 2C, bottom 2 panels). These results show that axon-related defects appear early, from the time axon structures tend to emerge and that the incidence of axon defects in KO embryos increases during development.

### Eip63E deficiency leads to other morphological defects in the CNS

To further evaluate the effects of Eip63E deficiency on neuronal cell bodies, we stained for the product of the elav (embryonic lethal abnormal visual system) gene which codes for an RNA-binding protein that is expressed exclusively in neurons (Robinow and White, 1991) and is required during the development and maintenance of the CNS (Jimenez and Campos-Ortega, 1987). ELAV expression pattern in Eip63E^81^ KO (GFP^-^) embryos revealed defects as early as stage 12 (Fig. 3A). ELAV staining is diffuse in the KO compared to GFP^+^ embryos of similar stages. Additionally, the morphology of neuromeres and hemisegments along the VNC of mutant embryos was disrupted and abnormal, making it difficult to distinguish midline neurons from the rest in several VNC segments of stage 12-13 KO embryos (Fig. 3A).

**Figure 3.**
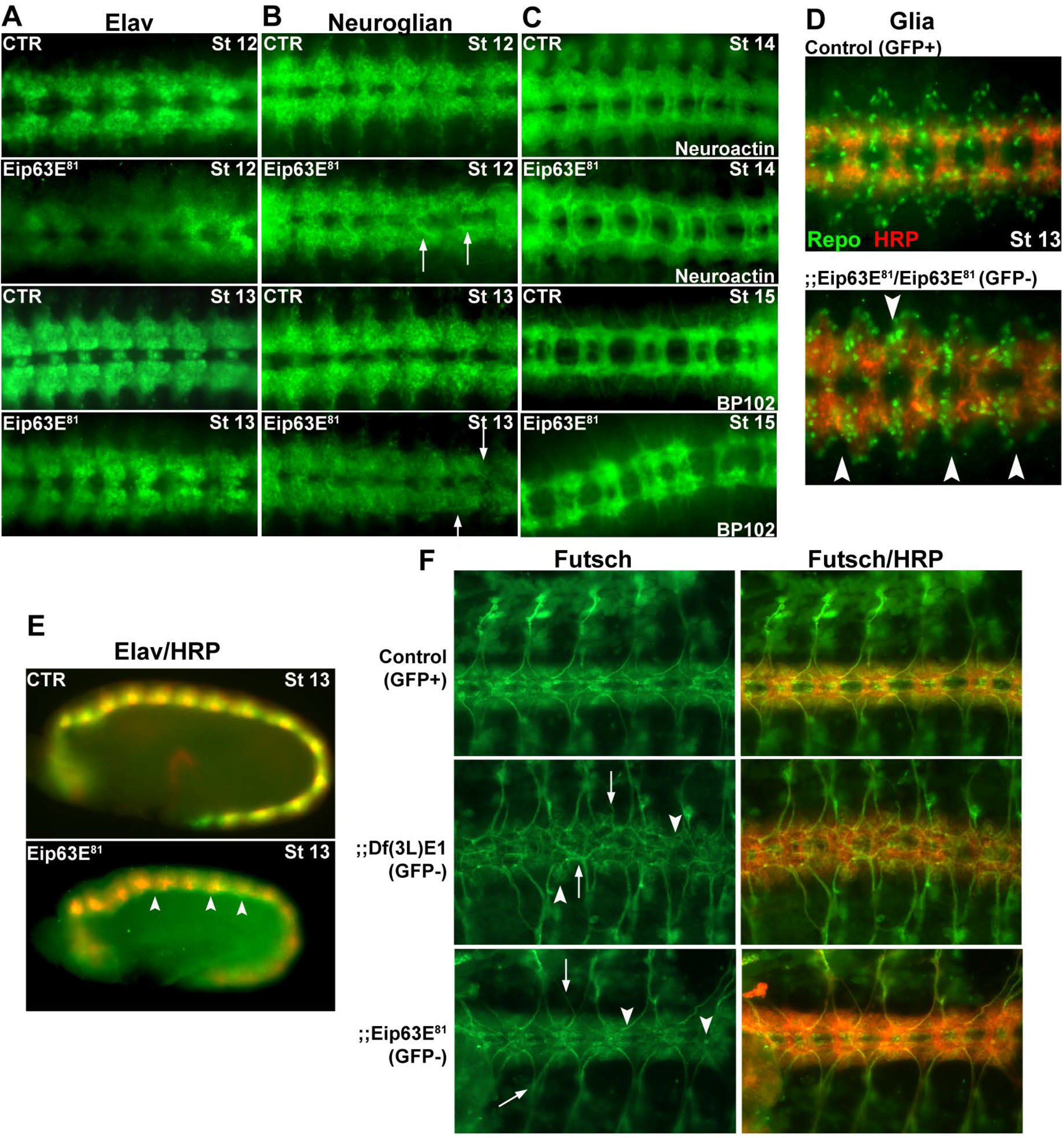
Eip63E deficiency leads to early segmentation defects. (A, B, C and D) Ventral views of whole mount embryos stage 12-15 shown head to the left. CTR show GFP+ embryos (wild-type and mutant heterozygous population) and Eip63E^81^ represent homozygous mutant embryos (GFP-). Immunostaining for Elav (A), Neuroglian (B), Neurotactin (C, top 2 panels) and BP102 (C, bottom 2 panels) showing axon defects (arrows). (D) Co-staining for glial populations (Repo) and axons (HRP) showing disorganized cell bodies (arrowheads). (E) Merged elav and HRP staining images (lateral view) of stage 12 embryos shown in panel A. Elav-positive cells appear to be dorsally deeper than normal (arrowheads), under the axon bundles. (F) VNC flat preparations of timed embryos (10-11h, stage 14) co-stained for Futsch and HRP are shown dorsal up, head to the left. Arrowheads point to some of the structures where neuronal organization is different from control embryos. Arrows point to examples of axon growth abnormalities

Lateral views of elav and HRP co-staining (Fig. 3E) reveal that Eip63E deficiency resulted in a disorganized neuronal cortex, since elav staining was detected dorsally underneath the axon scaffold. Similar abnormalities were observed using a different neuronal marker, Neuroglian (long isoform, BP-104), a cell-adhesion molecule expressed in a sub-population of neuronal cell bodies and axons (Hortsch et al., 1990). Malformation of neuromeres and hemisegments linked to axon pathfinding defects are observed in Eip63E^81^ KO embryos (Fig. 3B). The lack of symmetry between the hemisegments and the midline pattern of different neuromeres suggest a disorganization of neuronal cell bodies consistent with the observations for elav immunostaining. Disarray of neurons and axons was also observed when VNC flat preparations of timed stage 11 embryos were co-stained for Futsch and anti-HRP (Fig. 3F). Futsch is a microtubule-associated protein, necessary for axon and dendrite development in Drosophila (Hummel et al., 2000). The axonal bundles in Eip63E KO embryos (both Df(3L)E1 and Eip63E^81^ alleles) appear defasciculated when compared with controls (HRP staining) and defective hemisegments coincide with misplaced neurons (Futsch staining). Additionally, as can be seen with Futsch/HRP merged images, neuronal cell bodies are abnormally located dorsally to the axon scaffold in KO embryos, consistent with results from elav staining (Fig. 3E).

As in vertebrates, glial cells play a key role in neuronal development in Dmel, particularly in axon guidance (Chotard and Salecker, 2004; Freeman, 2006; Parker and Auld, 2004). Since the axon defects linked to Eip63E deficiency appear early in CNS development, we explored the effect of the deficiency on glial cells. The reverse-polarity (Repo) gene codes for a homeodomain protein that is expressed in all embryonic glial populations, except midline glial cells, from stage 9 glioblasts to mature glial cells (Xiong et al., 1994). Co-staining of glial cells using Repo and of axons using HRP antibody (Fig 3D), revealed disorganization of glial cell bodies in Eip63E^81^ KO embryos when compared to controls. Interestingly, those defects were much more severed (even with smaller cell population) for the Df(3L)E1 KO embryos (data not shown).

When we combined the penetrance associated to all these relevant neuronal and axonal markers along the VNC structures (Fig S1) it became apparent that Df(3L)E1 allele associated defects appear later in development with more sever outcomes (higher penetrance) than those associated with Eip63E^81^ allele. For both mutant alleles, axon defects were also accompanied by defects in neuronal cell body localization. This pointed to the possibility that axonal defects could be caused by mis-localization of neurons rather than a neuronal Eip63E requirement.

### Specific downregulation of Eip63E in neurons leads to axon defects

To further confirm that the axonal defects observed in the Eip63E KO mutants were indeed due to Eip63E deficiency during CNS development and rule out any background effects of the germ-line deletion, we performed tissue-time specific downregulation of Eip63E using the Gal4-UAS system (Duffy, 2002). Utilizing the UAS-RNAi fly lines for Eip63E developed by the Vienna Drosophila RNAi Centre, Eip63E was knocked down (KD) using drivers for all cells (ubiquitous expression, Act5-Gal4), neurons (elav-Gal4), glia (repo-Gal4), mesoderm/visceral muscles (mef2-Gal4), dorsal epidermis/ peripheral nervous system (PNS) (GawB109.69-Gal4) and digestive system/ longitudinal muscles (GawBtey5053A-Gal4) (Fig. 4). The Krüppel-GFP balancers used before would not be appropriate for these experiments, since they possess an internal Gal4-UAS system to regulate GFP expression that would interfere with our experiments. Therefore, lines were introduced to balancer chromosomes expressing ACT5GFP instead, which provided for GFP expression under the Actin A5c promoter (Reichhart and Ferrandon, 1998).

**Figure 4.**
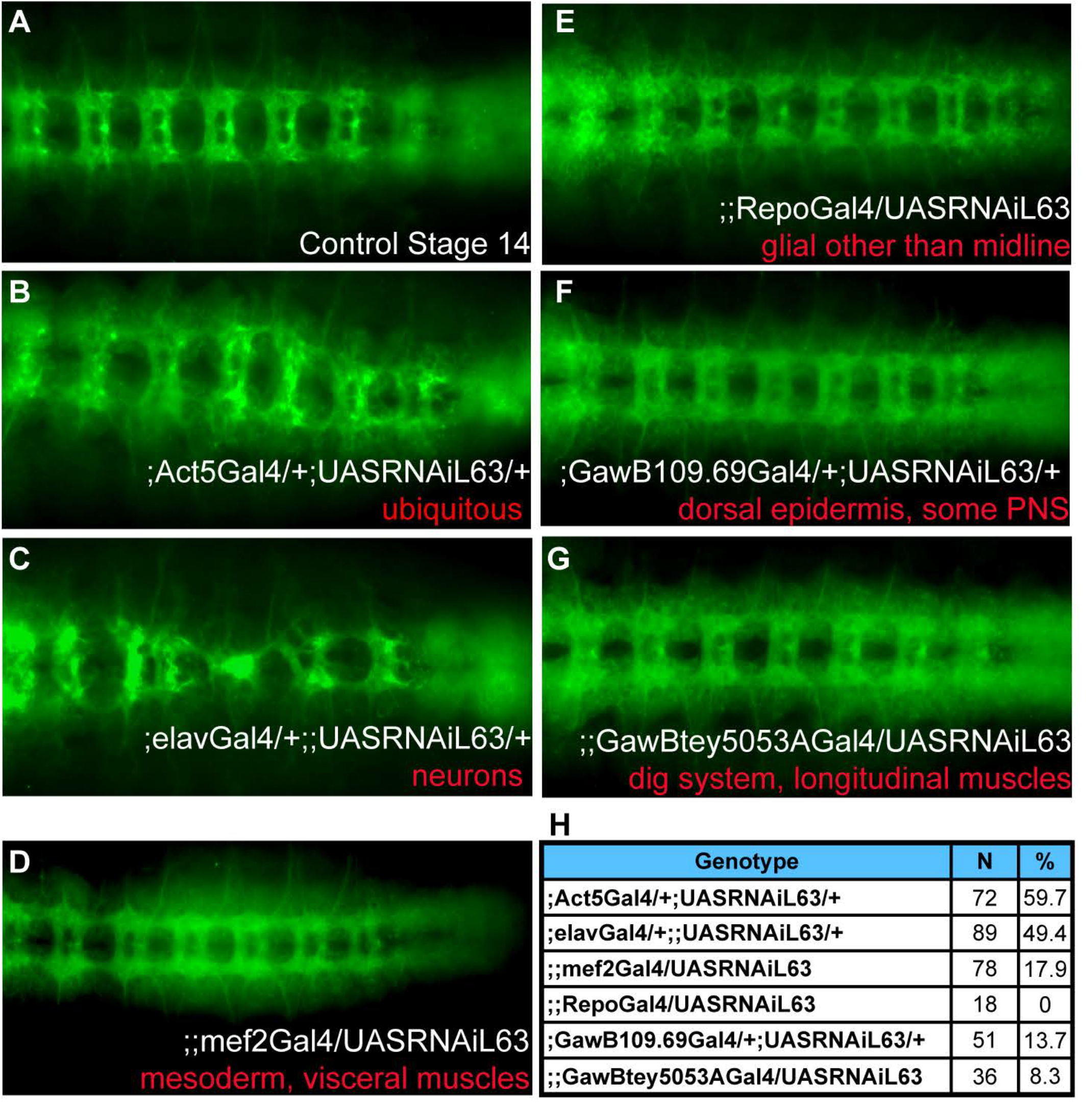
Only ubiquitous and neuronal downregulation of Eip63E leads to embryonic axon defects. Ventral views of stage 14 whole mount embryos shown head to the left. A UAS-Gal4 approach was used to drive Eip63E downregulation in specific tissues during embryonic development. Embryos negative for GFP (present in balancers for both the Gal4 and the UAS lines) was considered as experimental embryos vs the GFP positive of the same progeny. (A) A representative GFP+ embryo is shown. (B) Ubiquitous downregulation led to a replica of the germline deficiency lines, regarding CNS defective phenotype and developmental lethality (see text for details). (C) Neuronal-specific downregulation led to a robust and highly penetrant axonal phenotype. (D, E, F, G) Downregulation of Eip63E in other tested tissue (indicated in red) did not cause any detectable axonal defects. The number of embryos per genotype and the percentage of embryos per group showing axonal defects is summarized in (H).

As shown in Figure 4, ubiquitous down regulation of Eip63E resulted in axon defects and lethality similar to those observed with the germline deficient lines. Half of the embryonic population was expected to be control (no driver). Axon defects were observed in 59.7% of the embryonic population (Fig. 4B). This would suggest a 100% penetrance of defects for ubiquitous downregulation. Additionally, when embryos were left to complete the life cycle, all the flies that eclosed were positive for the balancer phenotypic marker, suggesting that (like the germ-line mutants), ubiquitous downregulation of Eip63E is developmentally lethal and corroborating the effectiveness of the RNAi construct used.

When Eip63E was specifically downregulated in neurons using the neuron-specific elav-Gal4 driver (Fig. 4C) VNC defects were observed in 49.4% of the embryonic population. Since half the embryonic population is expected to be control (no driver), these results suggest close to 100% penetrance of defects upon neuronal downregulation. These results are consistent with the notion that neuronal Eip63E function is required for axon development in Dmel. Intriguingly, when development was not interrupted for embryo collection, we observed an equivalent number of Cyo-positive and Cyo-negative flies eclosing (n=72 vs n=76). This suggests that although important in CNS development, Eip63E neuronal function alone does not play a role in developmental survival.

No noticeable defects were observed when Eip63E was specifically knocked down in glial, or any of the other tissues (Fig. 4 D to G). The total number of embryos per group and the percentage of embryos with observed axon defects are presented in Fig 4H. These results suggest that the functions of Eip63E in neurons, and possibly neuroblasts, are key to axon development in Drosophila, especially for the regulation of axon outgrowth and guidance.

### Drosophila Eip63E functionally interacts with Rho1

Previous reports have shown that CDK5 and PCTAIREs regulate axon development by functionally interacting with the Rho family of GTPases (Matsuda et al., 2014; Nikolic M, 1998; Rashid T, 2001). However, whether such an interaction exists between PFTK1/Eip63E and Rho GTPases is not known. We used the Dmel line expressing the dominant-negative Rho1^72F^ allele (Warner and Longmore, 2009), to explore this possibility. Homozygosity for the Rho1^72F^ mutant allele leads to 100% lethality and this germline was therefore maintained as a balanced heterozygous population using the Cyo-KrGFP balancer chromosome. When we characterized the embryonic population staining for BP102 marker, we detected that 97% of homozygous mutant embryos (stage 11-15, N=47) had acute VNC axonal defects. Moreover, 55% of the GFP^+^ (mix of heterozygous and WT) embryonic population was also defective (N=86), pointing to the dominant negative nature of the Rho1^72F^ mutant allele. One of those defective GFP^+^ embryos is shown in Figure 5C as reference. For the Eip63E^81^ line, since we never found a defective GFP^+^ embryo, we concluded that Eip63E^81^heterozygosity does not lead to axon defects (Fig. 1A, bottom left table and Fig. 1B bottom right table; Fig. 5B;).

**Figure 5.**
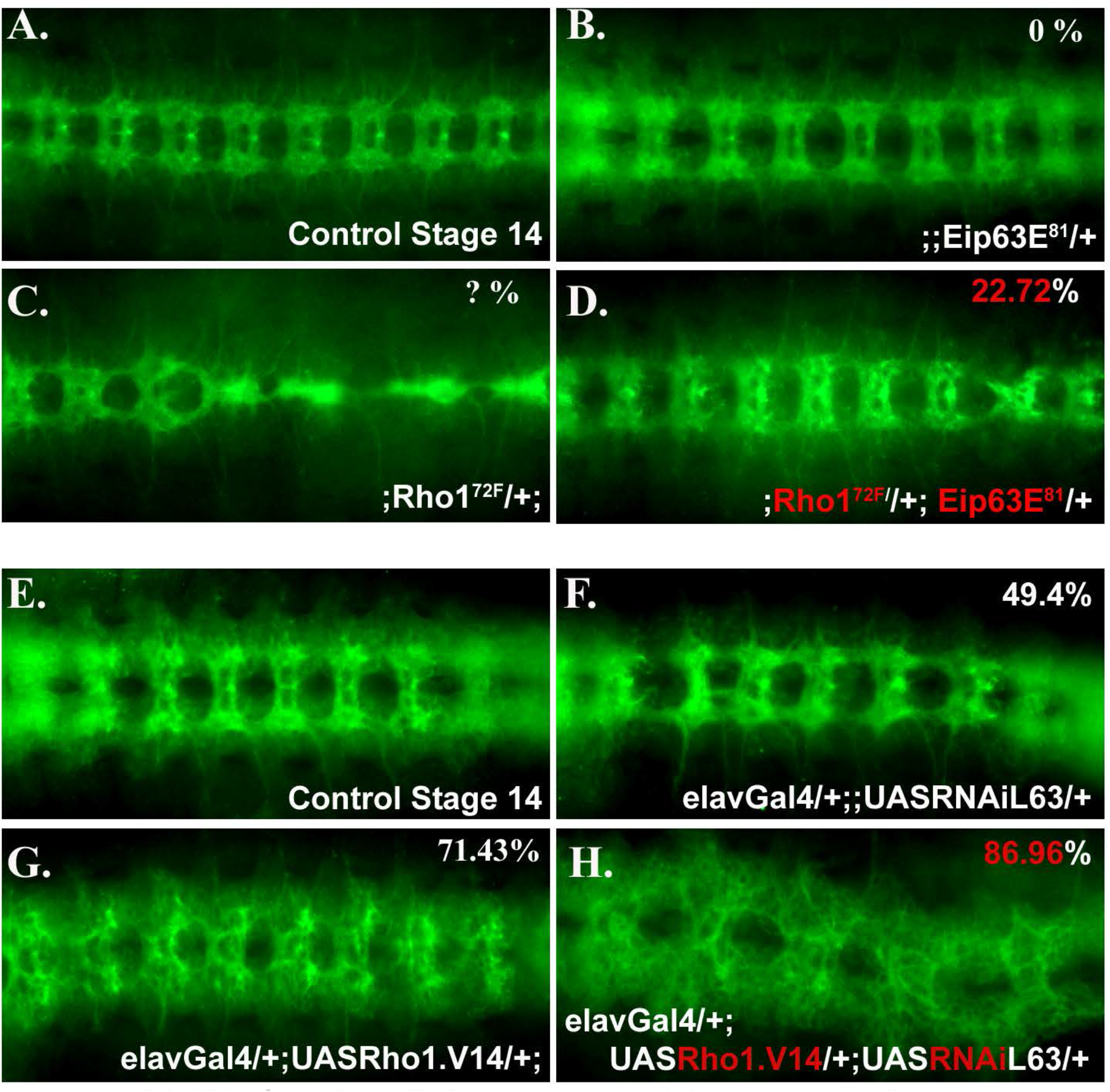
Eip63E functionally interacts with Rho1 to regulate axogenesis. Ventral views of stage 14 whole mount embryos stained for BP102 antigen are shown head to the left. Panels show representative images for each specified genotype, (A-D) Representative images of control embryos (A) single heterozygous Eip63E^81^ (B) or Rho1^72^F (C) from a cross between Eip63E^81^ KO line and Dominant-Negative Rho1 (Rho1^72^F). (B, C) Images of presumed genotype (see text for details). (D) Axon defects of intermediate severity in 22.7% of double-heterozygous mutants (GFP-). (E-H) UAS-elavGal4 approach to drive Eip63E downregulation and constitutive active Rho1 overexpression in neurons in two independent Dmel lines. Lines were then crossed to generate the indicated genotypes. (F, G) Heterozygous neuronal downregulation of Eip63E or Rho1 showing axon defects with specified penetrance. 86.96% of Double-heterozygous embryos (H) had axon defects showing features of both heterozygous controls. Penetrance values from stages 13-17, n as follows: B) >100, C) 86, D) 22, F) 89, G) 126, H) 69. n represents the total of screened embryos from several different pairings/collection pools.

We then crossed both the Eip63E^81^ and the Rho1^72F^ lines to create double heterozygous mutant embryos (GFP^-^). 22.72% of those embryos showed VNC defects that were of intermediate severity (Fig. 5D). Interestingly, that double heterozygosity also corrected Rho1^72^ lethality, since flies lacking the balancer phenotypic marker were present in the adult population when development was not interrupted for experimentation (multiple crosses, 155 live double-heterozygous adult flies out of 400 total N). These results suggests that there is indeed functional genomic interaction between the mutant alleles, suggesting that Eip63E and Rho1 are part of the same metabolic pathway in Dmel axogenesis.

We wanted to explore next if the same would happen for simultaneous Eip63E downregulation and Rho1 overactivation in neurons. Since these experiments would require using the UAS:Gal4 system, no possibility of genotyping embryos using GFP expression was possible, just like for the tissue/time specific expression experiments. We already observed that neuronal Eip63E downregulation (elavGal4:UASRNAiL63) leads to 49.4% of the embryonic population to be defective (50% expected to be expressing the allele) (Fig. 4C,H; Fig. 5F) and that it wouldn’t lead to any detectable lethality (76 vs 72 adult flies corresponding to absence and presence of balancer phenotypic marker). For Rho1 activation, we used a line expressing a constitutively active Rho1.V14 allele under a UAS promoter. When driven in neurons, 71.43% of the embryonic population showed to be defective (50% expected to express the allele) and no adult fly ever eclosed that didn’t show the balancer phenotypic marker (N=172) implying a 100% lethality.

We then created a double heterozygous line containing both the RNAiL63 and the Rho1.V14 alleles under UAS promoters and crossed it with a Gal4:elav line to drive neuronal expression. This resulted in 100% lethality since no adult fly ever eclosed that didn’t show the balancer phenotypic marker (n=117). Assessment of the embryonic population following BP102 staining revealed embryonic VNC axon defects in 89.96% of the population with axon phenotypes that resembled a combination of both the controls (Fig. 5H). We detected less axons like those observed in Rho1.V14 control (Fig. 5G) with acute defasciculation and misguidance, just like the RNAiL63 control (Fig. 5F). Since all the individual control phenotypes were present, we conclude that there is a lack of functional interaction between Eip63E deficiency and constitutively active Rho1.

We also examined any functional interaction between Eip63E and the other major members of the Rho family of GTPases, Rac1 and Cdc42 (Fig. S2). Results show that simultaneous Eip63E neuronal downregulation and activation of Rac1 and Cdc42 (constitutive active forms), but not downregulation (dominant negative forms), lead to rescue of the control phenotypes. This data suggests that in flies, the Rho family GTPases might be players in the same pathway as Eip63E regulating axonal development. Furthermore, based on our Rho1.V14 data, since Rho1 activation did not rescue the defects associated with heterozygous UASRNAiL63 embryos, our results also point to a role for Eip63E upstream of Rho GTPases, by regulating their activity or that of their regulators.

### PFTK1 downregulation and deficiency leads to faster-growing axons in murine cortical neurons

Previous research using murine experimental models has indicated that CDK5 inactivation leads to inhibition of axonal growth (Nikolic M, 1998; Nikolic M, 1996) and that CDK related protein kinase PCTAIRE1-3 negatively regulates neurite outgrowth (Cole A R, 2009). We investigated whether manipulation of PFTK1 activity had an impact on axon outgrowth in mammalian cells. Primary cortical cultures from CD1 mouse embryos were infected with adenovirus expressing GFP tagged-D228N PFTK1, GFP tagged-wild type PFTK1 or GFP control alone (Fig. 6A, B). Both at 12hr and at 24 hours after plating, a significant increase in axonal length was observed in cells expressing D228N but not the GFP control or WT PFTK1 (Fig. 6A, B). This suggests an increase in neurite length in vitro upon downregulation of PFTK1 function, contrary to that observed with the loss of CDK5 (Nikolic M, 1996).

**Figure 6.**
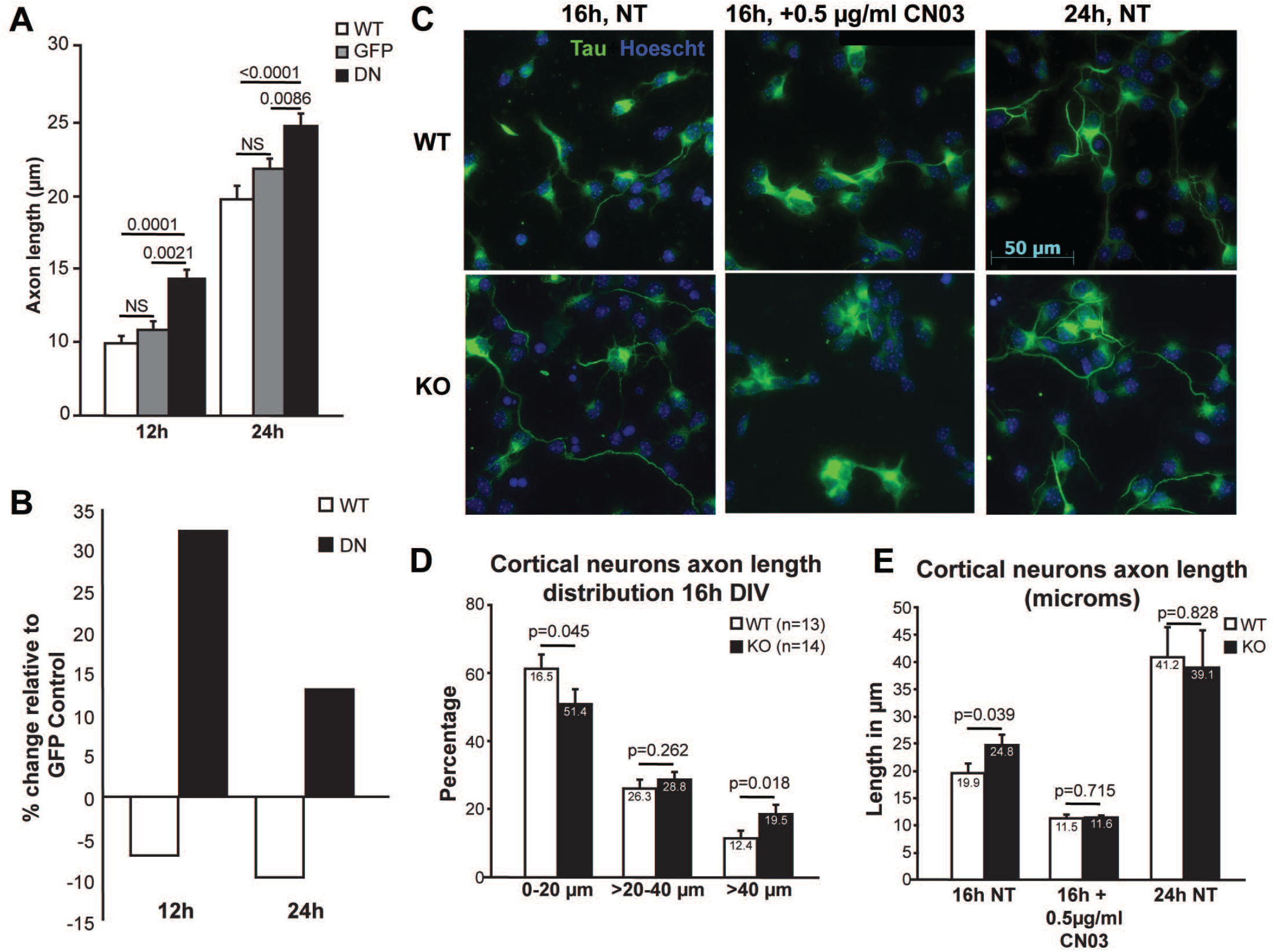
PFTK1 downregulation leads to faster-growing axons in murine cortical neurons. Primary cortical neurons derived from CD1 embryos (E 14-15) were infected with 100 MOI of adenoviruses expressing GFP alone, GFP and WT- PFTK1 or GFP and D228N- PFTK1. After 12 or 24 hours in culture, cells were fixed and stained with Hoechst 33258 (nuclei visualization). Axons of GFP+ neurons were measured using Image J. (A) Values are presented as averages ±SEM from 122-143 cells/treatment/timepoint. Statistics were done using two-way ANOVA followed by Tuckey’s post-hoc. (B). Same data as in (A) but presented as axon length relative to the GFP control. (C) Representative images of primary cortical neurons from E13-14 embryos from HET x HET crosses from WT and KO cultures fixed at the specified times and stained for Tau and Hoechst. (D) Axon length distribution per genotype from the indicated number of embryos (236-469 cells/embryo) at 16 hours in vitro. Each column represents the percentage of cells in the specified range of length for the indicated genotype. Non-treated (NT) cultures were also fixed after 16 and 24h in vitro. Graph represents the average axon length/genotype/treatment. t-test was used for statistical analysis whenever p value is specified. (E) 16h cultures were treated for the last 8h with 0.5ug/ml of RhoA activator (Cytoskeleton Inc. CN03). Graph represents the average axon length treatment.

We then generated a PFTK1 deficient mouse line to further examine the role of CDK14 in CNS development and maintenance. Details of the generation of the PFTK1 KO mouse line are presented in Fig. S3 and described in the methods section. Unlike PFTK1 deficiency in Drosophila, which leads to embryonic lethality, PFTK1 knockout (KO) mice are viable, develop normally and do not show any obvious morphological defects compared to the wild type littermates. Microscopic analysis of coronal sections of the adult mice brain on a mixed C57Bl6 background (3^rd^ generation) revealed no major abnormalities upon cresyl violet staining (Fig. S4A). Furthermore, no noticeable deviation from the expected Mendelian ratios (Fig. S4B) were observed in embryonic or postnatal populations, suggesting that PFTK1 loss alone, in mice, might not have the same effects observed with L63E deficiency in the fly regarding lethality.

Next, to investigate whether the axonal outgrowth effects of D228N overexpression could be replicated by PFTK1 deficiency, we studied populations of axons in primary cortical cultures from PFTK1 KO and WT littermate embryos (Fig 6C, D, E). Cultures were prepared from E13.5 - 14.5 mouse embryos, fixed at 16 and 24 hours after plating and stained for the microtubule marker Tau-1 and Hoescht. The axon length distribution at 16h in vitro (n=13-14 embryos/genotype, 236-469 cells/embryo) revealed a statistically significant increase in the number of axons longer than 40 µm in PFTK1 KO neurons and a decrease in shorter axons when compared to WT controls (Fig. 6C, 6D). Nevertheless, the significant difference between KO and WT cultures at 16h is no longer detected after 24h in vitro (Fig 6E), which would suggest that either the effect is transient (early axogenesis) or that axons had reached their maximum possible length in this model at the 24h mark. Together these results point to a role of PFTK1 in regulating axon outgrowth at least during early axogenesis.

### PFTK1 regulated axon outgrowth is dependent on RhoA activity

Based on the functional interaction of Eip63E and Rho GTPases, next we investigated whether manipulation of RhoA activity in PFTK1 KO neurons could rescue the effects on axonal outgrowth. Under the same settings as the experiments described above (same embryos), PFTK1 KO or WT neurons were treated with RhoA activator CN03, which blocks intrinsic, and GAP stimulated GTPase activity leading to constitutive activation of RhoA by robustly increasing the level of GTP bound RhoA (Flatau et al., 1997). Figure 6C and 6E show that treatment of the RhoA activator CN03 eliminates the difference between KO and WT cultures by equally decreasing the axon length of both population of cells (Fig. 6C and E). Consistent with our Dmel results, these data present a strong case for the role of RhoA activity downstream of PFTK1 and the specific involvement of PFTK1 mediated RhoA regulation in axonal outgrowth.

We then assessed whether PFTK1 levels influence RhoA expression. To be as relevant to the axon length experiments as possible, we decided to do the evaluation in extracts (total RNA and protein) obtained directly from cultured WT and KO cortical neurons (Fig. 7A-C). Reverse transcription polymerase chain reaction (RT-PCR) analysis showed no significant differences in RhoA mRNA level (Fig. 7A and C) in KO vs WT. Surprisingly, examination of the endogenous levels of RhoA protein revealed significantly higher levels of RhoA in PFTK1 KO cortical neurons compared to those of WT (Fig. 7B and C; Fig. S5). This was unexpected, since the genomic interaction experiments in Dmel and the CN03 experiment in cortical neurons were suggesting that PFTK1 deficiency might be leading to less Rho activity. That was also the expectation, since it is very well known that lower Rho activity leads to longer axons (Kranenburg et al., 1997). Then the question remained of weather the increased levels of RhoA protein in the KO cortex was actually equivalent to increased RhoA activity.

**Figure 7.**
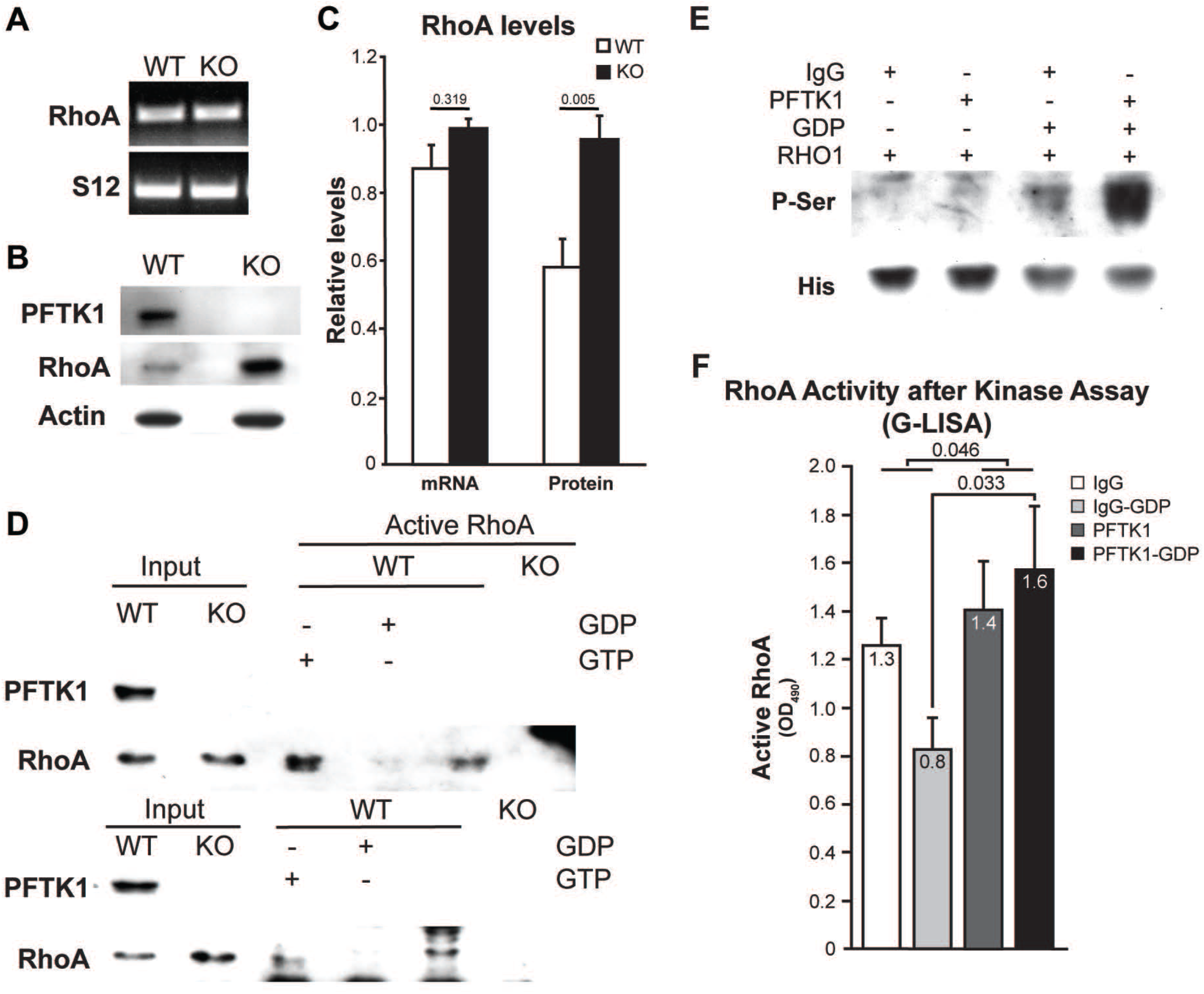
RhoA phosphorylation and activity is affected by PFTK1. Neuronal cultures from E13-14 embryos from HET x HET crosses were examined after 18-20h *in vitro*. Total RNA (A) or protein (B) was extracted followed by RT-PCR or western blot. Representative images from RT-PCR (A) and western blot (B) assays are shown. (C) Bar graph represents the average band intensity from 3 KO and 6 WT for mRNA and 7 WT and KO each for protein. t-test was used for statistical analysis. D) Assessment of RhoA activity using an IP-based assay. Neuronal extracts were incubated with Rhotekin PBD beads to pull down active RhoA. WT extract preincubated with GTPγS (positive) or GDP (negative) were used as controls. IP was followed by western blotting for RhoA. Two representative assays of four independent assays are shown. E) *In vitro* Kinase assay. IPed protein (either with IgG control or PFTK1) from WT cortical neuronal culture extracts was used as source of kinase and incubated with human recombinant RhoA (His-tagged) with or without GDP. Results were analyzed for phospho-serine signal. F) RhoA activity assay following *in vitro* kinase assay. Kinase reactions were analyzed by G-LISA to test for RhoA activity. Bar graph represents mean values ±SEM. Two-way ANOVA analysis of grouped data and Sidak test for multiple comparisons are indicated on bar graph. RHO1: His-RhoA human recombinant protein.

Of note, no significant difference in Rac1 or Cdc42 levels were observed between KO and WT cortical neurons (Fig. S5). Nevertheless, preliminary data from performing the same studies using the rest of the brain (no cortex or hippocampus) of the same WT and KO littermate embryos used for cortical cultures, suggest that the expression profiles might be inverted. In those extracts, Cdc42 and Rac1 seem to be upregulated in the KO samples vs WT (data not shown).

To answer the pending question, we assessed RhoA endogenous GTPase activity in PFTK1 KO vs WT cortical neurons (Fig. 7D) Briefly, cell extracts were prepared from cortical cultures of PFTK1 wild type and KO littermates at E13.5-14.5 and then incubated with Rho-binding domain (RBD) of Rhotekin RBD beads, which pulls down the endogenous active form of RhoA from the cell lysate. Active RhoA levels were examined by western blotting with RhoA antibody. The detected signal would correlate with RhoA activity and so we expected that the KO signal would be higher than the WT one. Interestingly, despite RhoA protein upregulation in PFTK1 KO mice, we did not detect any RhoA activity in the KO extracts with this technique (Fig. 7D). This suggested that the increased RhoA protein levels in the KO cortex is likely an attempt to compensate for lack or dramatically decreased RhoA activity.

### RhoA phosphorylation and activity is affected by PFTK1

So far, all our results pointed to a close functional interaction between Eip63E/PFTK1 and Rho1/RhoA but without clarifying if such an interaction was direct. To address this, we decided to test if RhoA would be a substrate for PFTK1 (Fig. 7E). Since no recombinant PFTK1 exists yet and it is not clear which and how many cyclins/ coactivators it requires to become active, we were left with the alternative of using cortical neurons extracts from E13-14 WT embryos as source of our kinase. Immunoprecipitation would render PFTK1 (presumably attached to all relevant co-factors). Optimization of the process was done using different antibodies, clearing and incubation times to obtain the cleanest and most PFTK1 enriched sample possible (data not shown). As control, IgG immunoprecipitated samples were used.

Using recombinant His-tagged RhoA as substrate, we then proceeded to optimize an in-vitro Kinase assay in the presence or absence of immunoprecipitated PFTK1, followed by Western blot analysis for the resulting reactions using a phosphoserine antibody (Fig. 7E). Noticing the inconsistency of preliminary results that only used the recombinant, we then tested if adding GDP to the reaction mix would stabilize RhoA as a substrate.

Figure 7E shows one representative experiment. In all replicas of the experiment, a noticeable increase in serine phosphorylation of recombinant RhoA was observed only in the presence of PFTK1 and when GDP was externally added to the reaction (Fig. 7E, last lane to the right). These results suggest that recombinant GDP-RhoA is a direct in vitro substrate of PFTK1.

It was important then to determine whether PFTK1 phosphorylation of GDP-bound RhoA had any effect on RhoA activity, since previous reports have shown that depending on the phosphorylation site, RhoA can be activated or deactivated (Ellerbroek et al., 2003; Forget et al., 2002; Tong et al., 2016). To this effect, we submitted all reactions’ products of the in vitro Kinase Assay described in the previous experiment (Fig. 7E) to a G-LISA reaction to test for RhoA activity (instead of using then for western blot). This method, being semiquantitative and more sensitive than the IP approach used before (Fig. 7D), revealed a significant increase in RhoA activity as a function of PFTK1 compared to IgG control. Additionally, a significant increase in RhoA activity was observed when GDP bound RhoA was incubated with PFTK1. These results suggest that GDP-RhoA is not only a substrate of PFTK1 but also that this phosphorylation results in RhoA activation (Fig. 7F). Taken together, these results point to a yet unidentified role of PFTK1 in regulating RhoA activity by phosphorylating and thereby activating GDP bound RhoA.

## Discussion

The proper development of the CNS requires tight coordination of several important processes including axon development. This is a complex process involving a broad range of factors that are present in the neuronal microenvironment. However, the precise role of PFTK1 in neurite morphogenesis and implications in CNS development has not been fully characterized. Here we present evidence that PFTK1 (Eip63E in Drosophila), is involved in this process and mediates its action on axon development by regulating its downstream effector via its kinase activity.

Screening of Eip63E deficient embryos from two different and independent mutant alleles consistently show defects in VNC structures. Results (Fig. 1A, bottom panel) showing that the phenotype of the trans-heterozygous embryos is less severe than that of the Df(3L)E1, which contains a larger deletion with at least 2 other background mutations, does indeed suggest that these background mutations have a role in the homozygous Df(3L)E1 phenotypes. Nevertheless, since the only homozygously deleted gene in the trans-heterozygous embryos is Eip63E, our results support the interpretation that the axonal phenotype is linked to Eip63E. This was further confirmed when only neuronal downregulation of Eip63E led to the same VNC defects than those in the mutant germlines. These axon defects occur along neuronal and, to a lesser extent, along glial cell body disorganization. The penetrance values of neuron-related defects are comparable to those of axonal defects already from stage 10 onward (Fig. S1). This does not seem to be the case for glial defects. No consistent glial perturbations were noticed between the two Eip63E mutant alleles. Taken together, these observations suggest that the axon defects are related to neuronal, rather than glial-related Eip63E functions.

More than one of the CNS markers used to characterize the VNC phenotype of Eip63E KO embryos, point to a potential defect in biological processes that regulate or mediate cell identity specification and/or neuronal differentiation processes (Fig. 2B and Fig. 3). These defects could be described in general as an abnormal cell cortex laminar organization along the VNC of mutant embryos, a process that is highly dependent on the timing and pattern of progenitors’ birth and divisions (Isshiki et al., 2001). It is possible that defects in Eip63E mutants are related to temporal control of new neurons and/or their differentiation. Indeed, it has been shown before that the absence of cell cycle progression or cytokinesis in neuroblasts leads to abnormal expression pattern of hunchback (Hb) (Grosskortenhaus et al., 2005), one of the progenitor temporal transcription factors that regulate first-born cell fates in Drosophila (Isshiki *et al*., 2001). Previous reports show that Eip63E has a role in cell cycle regulation, since its downregulation in embryonic cell cultures by RNAi leads to low mitotic index and therefore larger G2 cells (Bettencourt-Dias et al., 2004). There is also evidence that Eip63E interacts with cyclinY to phosphorylate the low-density lipoprotein receptor-related 6 (LRP6; ‘arrow’ in Drosophila) and mediate Wnt signaling (Davidson et al., 2009); a pathway broadly recognized as essential for regulating progenitor proliferation, cell lineage decisions and neuronal differentiation in insects and vertebrates (Grigoryan et al., 2008; Zechner et al., 2003).

In addition to this early potential function for Eip63E, our data strongly support a later role for this kinase in neurons to regulate axon development. It is evident that early deregulation of differentiation timing would set a precedent for axon defects, since for example, pioneer neurons and axons are needed for correct pathfinding and fasciculation of more mature axon pathways (Sanchez-Soriano and Prokop, 2005). A key concern for the interpretation of these results is whether axogenesis disregulation is the primary cause of the axonal defects and not any of the other described phenotypes observed as a result of Eip63E deficiency. The observation that Eip63E downregulation in elav-driven cells (differentiating neurons (Yao and White, 1994)) is the only tissue specific downregulation that resulted in VNC defects (Fig. 4C) presents a strong argument for the role of Eip63E in neuronal function. Consistent with this observation, downregulation of Pftk1 in vitro in murine embryonic cortical neurons also resulted in accelerated axon outgrowth. Interestingly in the mouse model, the other observed phenotypes associated with Eip63E deficiency in the fly model, including lethality, was not observed. This suggests that Eip63E could have key functions in other biological processes outside of the CNS in the fly model that might be taken over by other Cdks in mice.

A hypothesis driven search for functional interactors of Eip63E activity resulted in the identification of RhoA as a potential target. RhoA belongs to the Rho subfamily of GTPases within the Ras superfamily, which also include Cdc42 and Rac1. The family which now includes 20 total members (Hodge and Ridley, 2016) is involved in diverse cellular functions such as the regulation of cytoskeletal rearrangements, cellular motility, cell polarity, axon guidance, vesicle trafficking and the cell cycle (Heasman and Ridley, 2008). Most of the Rho family of proteins function as switches that undergo conformational change between the inactive GTP-bound and the active GTP-bound state. Guanine nucleotide exchange factors (GEFs) activate these Rho GTPases by regulating the dissociation of GDP and the subsequent uptake of GTP from cytosol whereas GTPase-activating proteins (GAPs) facilitate their inactivation by increasing the rate of hydrolysis of GTP. Inactive Rho GTPases in the cytosol are often bound to guanine nucleotide dissociation inhibitors (GDIs) which prevent the GTPases from interacting with their Rho-GEFs at the plasma membrane (Garcia-Mata et al., 2011), providing an additional layer of regulation. The role of RhoA in neuronal morphogenesis has been long implicated and has become clearer over the past several years. Although controversial, with initial overexpression studies suggesting inhibition of RhoA activity promoting neurite outgrowth (Jin and Strittmatter, 1997; Kozma et al., 1997; Kranenburg *et al*., 1997), several others (Kuhn et al., 1999; Threadgill et al., 1997; Zipkin et al., 1997) demonstrate that RhoA activity positively affects neurite initiation and extension resulting in longer axons. Our results side with the former, pointing to a role for PFTK1 in regulating activation of RhoA leading to more controlled axogenesis. Both expression of a dominant negative form of RhoA and PFTK1 deficiency, resulting in reduced RhoA activity, leads to uncontrolled initiation of axon outgrowth and hence longer neurites.

When Rho1 activity was overexpressed in specific neuronal populations with Eip63E deficiency, VNC defects in axonal defasciculation and misguidance appeared to be additive as both features of Eip63E deficiency (Fig. 5F) and Rho1 constitutive active overexpression (Fig. 5G) could be observed (Fig. 5H). In contrast, simultaneous expression of the constitutive active forms of Rac1 (Fig. S4D) and Cdc42 (Fig. S4J) and Eip63E downregulation in neurons resulted in an intermediate phenotype, implying that Eip63E may be regulating upstream factors that activate Rac1 and Cdc42 and may need further investigations. Intriguingly, in mice cortical neurons, no change in Rac1 or Cdc42 expression levels were observed (Fig. S5). One possible explanation for this observed difference between the flies and mouse could be compensation by PFTK2 and PCTAIREs in mice. In flies, in the absence of these homologs, Eip63E emerges to have a dominant role in CNS development and regulation of the activity of Rho family of GTPases. Nevertheless, based on our results with Rho1 activity in flies, we conclude that PFTK1 functions upstream of Rho1, and PFTK1 deficiency results in axonal defects due to perturbations in Rho1 activity. CDK5 and PCTAIRE, two close family members of PFTK1 have been implicated in axon outgrowth (Nikolic M, 1998; Nikolic M, 1996). In our experiments, a noticeable change in axon lengths at earlier time-points (Fig. 6) might indicate that PFTK1 is more important in regulating the initiation of axon outgrowth rather than being involved in axon outgrowth. Interestingly, the percent of neurons with longer axons are higher at 16 hours in comparison to 24 hours (Fig 6D). This indicates that over time, the rate of axonal outgrowth evens out and the effect is lost, suggesting the participation of other downstream factors that regulate axon elongation. Note that this change in axon lengths is not due to generalized effect on the health of the cell as the expression of the dominant negative PFTK1 had no effect on the survival of neurons (Fig. S6).

The lack of any obvious gross CNS phenotype in PFTK1 deficient mice compared to those observed in flies, although intriguing, could be due, in part, to the presence of other PFTK1 related kinases. Flies merely possess the PFTK1 homologue Eip63E but not PFTK2 and PCTAIRE kinases. In comparison, mammals contain both PFTK1 and 2 that show 68 % homology, as well as PCTAIRE 1-3, which are 61 % homologous to PFTK1 and are all highly expressed in the mouse brain (Cole A R, 2009; Hirose T, 2000; Mikolcevic P, 2012; Okuda T, 1992). The presence of these highly similar genes in the mammalian system could compensate for the lack of PFTK1 and result in a lower level of abnormalities in the KO mice. However, the full extent of the effect of PFTK1 deficiency on brain morphology would require a closer examination of specific brain structures. Although the lack of gross developmental defects is not inconsistent with effects on axon length, future analysis of the PFTK1 mice would require a more careful examination.

We observe a noticeable increase in RhoA protein level in PFTK1 KO cortical neurons (Fig. 7B and C) despite no detectable increase in mRNA levels suggesting a posttranslational regulation of RhoA levels. It could be speculated that PFTK1 also plays a role in regulating cellular RhoA levels by regulating RhoA phosphorylation and ubiquitination. In this regard, previous reports have identified specific E3 ligases, Cullin-3, that targets GDP-bound RhoA for degradation (Chen et al., 2009) and further studies may provide clues to the role of PFTK1 in this pathway. Intriguingly, this increase in RhoA level did not translate to more active RhoA but rather a noticeable decrease in detectable active RhoA in PFTK1 KO neurons (Fig. 7D) suggesting that PFTK1 deficient neurons had reduced GTP-bound active RhoA. Additionally, PFTK1 deficiency also resulted in perturbation in axon length of cultured cortical neurons. These results are in line with previous studies that show that reduced RhoA activity leads to longer axons (Sebok A, 1999). Combined with the observations that pharmacological activation of RhoA resulted in the loss of the observed difference between PFTK1 WT and KO neurons (Fig. 6E), our data strongly supports a model where PFTK1 regulates RhoA activation which in turn mediates the initiation of axogenesis.

Observations via an in vitro kinase assay lend further support to this model (Fig. 7E). Serine phosphorylation of recombinant RhoA by PFTK1 in the presence of GDP indicates a preference for GDP-bound RhoA by PFTK1. Downstream targets or substrates of PFTK1, other than cyclin Y, that may play prominent role in its function in neurodevelopment, are not currently known. However, our data clearly establishes RhoA as a downstream effector of PFTK1 and identifies regulation of GDP-bound RhoA phosphorylation and activation by PFTK1 as a critical factor in PFTK1 mediated axogenesis. It is intriguing that addition of GDP to purified recombinant RhoA would have any effect on RhoA activity as purified RhoA is expected to already be in the GDP-bound state due to RhoA’s intrinsic GTPases activity. It should be noted that adding GDP to recombinant RhoA in the absence of Pftk1 resulted in reduced RhoA activity (Fig. 7F) suggesting a potential exchange of nucleotide under the conditions employed. Given that the addition of GDP resulted in increased phosphorylation of RhoA, it should be noted that regulation of Cdk family of proteins, of which Pftk1 is also a member, by GDP is not known in published literature.

Accumulating evidence suggests the importance of the regulation of Rho GTPases phosphorylation in its function (Chang et al., 2011; Tong *et al*., 2016). In this regard, it will be critical in future studies to address potential phosphorylation sites on RhoA, regulated directly or indirectly by PFTK1. All our attempts to address those questions did not render consistent results, as the approaches (GC-MS, targeted RhoA mutagenesis) were impeded by the complexity of obtaining reliable immunoprecipitated PFTK1 samples. Preliminary data suggest that the PFTK1 targeted site is conformational in nature, but more exhaustive experiments need to be done and more powerful approached applied to be able to answer this question.

Based on our evidence, it is also difficult to rule out the regulation of other upstream factors by PFTK1 that may mediate RhoA activity. This would add a further mode of regulation of RhoA function by PFTK1. Taken together, the evidence presented here proposes a working model for PFTK1 in mediating neuronal polarization and axon growth via RhoA regulation, pointing to the only known activating phosphorylation RhoA event with a role in axogenesis.

## Supplemental figure legends

**Figure S1. Eip63E deficiency leads to concerted defects in axons and neurons of the Drosophila VNC**.

The combined penetrance of markers relevant to the same VNC structures are presented as one. Axon-associated penetrance reflects those calculated using HRP, BP-102 and Fasciclin-II antibodies. Neuron-related values include Neuroglian, Elav and Futsch-revealed penetrance. Penetrance was calculated as the % of defective KO embryos from the total KO embryos screened. Both deficient lines show the same nature of defects. Although it seems that defects associated to the Df(3L)E1 allele are more acute and appear later in development, Eip63E81 shows a robust axonal phenotype. ND: Not determined.

**Figure S2. Eip63E functionally interacts with Rac1 and Cdc42 in D. melanogaster to regulate axogenesis**.

Ventral views of stage 14-15 whole mount embryos stained for BP102 antigen are shown head to the left. Panels show representative images for each specified genotype. An UAS-elavGal4 approach was used to drive Eip63E downregulation and either constitutively active or dominant negative forms of Rac1 or Cdc42 overexpression in neurons of independent D mel. lines. Fly genotype following crosses are indicated. (A-F) Neuronal functional interaction between Eip63E and Rac1. Only the combination of Eip63E deficiency and constitutively active Rac1 (UAS Rac1 V12; D), but not DN-Rac1 (UAS Rac1 N17; F), leads to an intermediate phenotype. (G-L) Functional interaction between Eip63E and Cdc42. Ventral views of the indicated genotypes of crosses between Eip63E and Cdc42 constitutive active form (UAS Cdc42 V12) or the dominant negative Cdc42 (UAS Cdc42 N17) are presented.

**Figure S3. Strategy for generation of PFTK1 KO murine line**.

The mouse was designed and developed by the Texas A&M Institute for Genomic Medicine (TIGM) (A) Strategy. Exon 6 (from nucleotide 18-Exon6 to nucleotide l-Exon7) of the Cdk.14 gene at chromosomes (4,803,391-5,380,19 7) was replaced for an IRES/bGeo/Poly A cassette by homologous recombination. (B) Southern Blot screening of clones. (C). PCR strategy for genotyping. Bottom panel shows representative gel image with samples from the three possible genotypes.

**Figure S4. PFTK1 deficiency does not appear to cause any gross brain defects or lethality**.

Brains from three-month-old mice (3rd generation backcrossed) were collected, perfused and fixed for sectioning. 14µm sections were collected from cryopreserved samples, starting from the cortex at bregma: ∼ 2.710 mm up through the 4th ventricle at bregma: ∼-4.20 mm. Sections were then stained with cresyl violet to examine the gross anatomy of the brain. No abnormality was detected for KO mice (n=3/genotype). Representative sections from bregma 1.98, 1.70, 0.74, 1.00, 1.64, and 2.75 mm are presented. (B). Embryos or mice (=> 6th generation backcrossed) from HET x HET crosses were genotyped and incidence of the three possible outcomes is presented as % of the total n. No gross deviation from the expected 25:50:25 Mendelian ratio was detected.

**Figure S5. Differential expression of Rho GTPases in brains of Wt vs KO E13-14 murine embryos**.

Only RhoA is dysregulated in cortical neurons. Basal levels of RhoA, Rac1 and Cdc42 protein in WT and Pftk1-defficient cortical neurons. Primary cortical neurons from E13-14 embryos from HET x HET crosses were dissected and plated at 0.65-0.75×106 cells/ml. After 18-20h in vitro, proteins were extracted followed by western blot. Top: Representative images from western bot are shown. Bottom, levels were assessed by densitometry using Image J and Rho GTPases signals were standardized vs loading control (Actin). Each column represents the average from 3 embryos/genotype +SEM. Significant difference between WT and KO neurons was only found for RhoA levels. Each column represents the average from n embryos/genotype +SEM. Significant difference between WT and KO neurons was only found for Cdc42 levels. t-test was used for statistical analysis whenever a p value is specified

**Figure S6. Manipulation of PFTK1 levels has no impact on neuronal survival**.

Primary cortical neurons derived from CD1 embryos (E 14-15) were infected with 100 MOI of adenoviruses expressing GFP alone, GFP and WT- PFTK1 or GFP and D228N- PFTK1. After 12 or 24 hours in culture, cells were fixed and stained with Hoechst 33258 (nuclei visualization). nuclear integrity of GFP+ neurons was used as a criterion for vitality. Survival values represent the % of healthy nuclei of the total GFP+ cells. Table is showing average +SEM from three independent experiments. Graph shows the same data but presented as value relative to the GFP control.

## Declarations

### Ethics approval and consent to participate

Not applicable

### Consent for publication

Not applicable

### Lead contact

All requests for further information, reagents and resources should be directed to the lead contact, Alvin Joselin.

### Availability of Data and Materials

Source data are provided with this paper. All other data supporting the findings of this study are available from the corresponding author on reasonable request.

### Author contributions

YRG and DSP conceived the project. YRG and FK designed the experiments. YRG, FK, PJ, SW, DQ, and DH performed experiments. PA designed, cloned and supplied the PFTAIRE constructs. LSA produced the adenoviruses. MS provided some of the fly lines and technical advice on fly work. YRG and AJ wrote and technically edited the manuscript. DSP and AJ are responsible for general supervision of research and administrative support.

## Materials and Methods

### Drosophila strains

Flies were kept under standard conditions. Stocks are described in FlyBase (http://flybase.bio.indiana.edu). The following stocks were obtained from the Bloomington stock centre: W1118, Eip6381/TM6tb, Df(3L)E1/TM6, Rho1^72F^, green balancer flies (L2Pin1/CyokrGFP, D*gl3/TM3krGFP, In(2LR)nocSco,b1/CyoActGFP and Sb/ TM3ActGFP,Ser) and Gal4 driver lines (elav::Gal4, Repo::Gal4/TM3,Sb, elav::Gal4;UAS::Dcr2.D and UAS::Dcr2.D;Act5::Gal4/Cyo). Three independent RNAi Drosophila lines were obtained from the Vienna Drosophila Rnai Center (UAS::RNAiL63/Cyo, UAS::RNAiL63 and UAS::RNAiL63/TM3,Sb). For the purpose of mutant embryos characterization, the TM6 balancer in both Eip63E deficient lines (Stowers *et al*., 2000) was replaced with the TM3.Kruppell-GFP balancer chromosome (Casso *et al*., 2000). To perform Eip63E specific downregulation, the Gal4:UAS system was used (Duffy, 2002). Balancers in all UAS and Gal4 lines involved in the crosses strategies were changed for the Actin5c-GFP balancer chromosomes (Cyo or TM3 as needed) (Reichhart and Ferrandon, 1998).

### Drosophila Immunohistochemistry

Drosophila embryos from Eip63E mutant lines were collected for 2 hours on grape juice-agar medium (Ashburner, 1989), and allowed to carry on development to embryonic stages 6-15 (Campos-Ortega and Hartenstein, 1985), in batches including only 2-3 stages each, by further incubation at 25°C. This developmental window includes all events of embryonic CNS development, from anlage formation to differentiation and axogenesis (Weigmann *et al*., 2003). Whole mount embryos were prepared following standard procedures (Harlow and Lane, 2006). Embryos were dechorionated with 50% bleach, followed by fixation for 20 minutes in heptane saturated with 4% Paraformaldehyde in 136 mM NaCl, 2.68 mM KCl, 10.14 mM Na2HPO4, 1.76 mM KH2PO4, pH 7.4, (PBS). Vitelline membrane was removed by vigorously shaking the fixed embryos in methanol, followed by further methanol washes. Embryos were re-hydrated using PBS-0.1% Triton X-100 (PBT) and then blocked for 20 minutes using 20% Normal Goat Serum (NGS) (GE Healthcare, Amersham RPN410) in PBT. Fluorescence immuno co-staining was performed against GFP (to amplify the presence of the balancer embryonic marker), HRP and a CNS marker. Primary antibodies were as follows: mouse anti-GFP (JL-8, Living colors, 1/100), Cy3-anti-HRP (1/25), mouse BP102 (1/10), mouse anti-En/Inv (4D9, 1/10), mouse anti-single-minded (1/5), mouse anti-Futsch (22C10, 1/10), mouse anti-Neuroglian (BP-104, 1/10), mouse anti-elav (9F8A9, 1/14) and mouse anti-Repo (8D12, 1/10). All monoclonal antibodies against CNS markers were obtained from the Developmental Studies Hybridoma Bank developed under the auspices of the NICHD and maintained by The University of Iowa, Department of Biology, Iowa City, IA 52242. Such antibodies were developed by C. Goodman, S. Crews, S. Benzer and G.M. Rubin. Since the anti-HRP antibody was Cy3-conjugated, only Alexa 488-Goat anti-mouse (1/100) was used as secondary antibody, after a second blocking step. Stained embryos were then directly mounted in Vectashield (Vector laboratories, H-1000) and visualized under a fluorescent microscope.

Embryo dissections were performed essentially as described (Benveniste et al., 1998). 7-13 hours timed embryos were transferred to a piece of double-sided tape mounted on a polylysine-covered microscope slide, and then rolled using forceps to mechanically remove the chorions. Embryos were then covered with PBS and examined under fluorescence light for GFP expression. Absence of GFP expression would indicate an Eip63E KO. After genotyping and fine determination of developmental stage based on morphological features, a dorsal incision was made in the vitelline membrane of each with a sharpened tungsten needle (Small Parts Inc TW-006-60). Each embryo was then removed from the membrane and placed ventral side down directly on the polylysine-covered surface of the slide, while still submerged in the PBS. Using the tungsten needle, an incision was made longitudinally along the dorsal surface of the embryo and the body walls were pressed down onto the glass. The gut and remaining yolk sac were then removed, resulting in a flat embryonic preparation. Dissected embryos were then fixed for 30 minutes at room temperature in 4% paraformaldehyde in PBT. One-hour blocking was performed using 10% NGS in PBT, followed by antibody staining. Fluorescence co-immunostaining was performed against HRP and a CNS marker. Same antibodies mentioned above were used but at different concentrations: 1/50 for primary antibodies and 1/300 for secondary. Preparations were mounted using Fluoromount™ mounting media (Sigma F4680), sealed with clear nail polish and visualized under a fluorescent microscope.

### Transgenic Mice Systems

All experimental animal studies were conducted in accordance with the guidelines of the University of Ottawa Animal Care Committee and conformed to the guidelines set forth by the Canadian Council on Animal Care and Canadian Institutes of Health Research.

### Generation and analysis of PFTK1 Transgenic (knockout) Mice

PFTK1 heterozygote mice were generated by the Gene Targeting and Transgenic Facility of Texas A&M Institute for Genomic Medicine (TIGM), on mixed (129/SvEvBrd x C57BL/6) background. To generate the mutant allele, PFTK1 Gene (MGI: 894318) and the transcript sequence of PFTK1 or (Cdk14-002 ENSMUST00000030763) located on chromosome5 (4,803,391-5,380,197) (reverse strand) was targeted. The intron-exon structure of the PFTK1 gene obtained by genomic mapping is illustrated in Fig. S3A. PFTK1 heterozygotes were generated by replacing the majority of exon 6 of the Mus Musculus PFTK1 transcript (ENSMUST00000030763) starting from nucleotide 18 with the neomycin (Neo) resistance cassette via homologous recombination. The resulting insertion in the PFTK1 gene causes a frame shift altering the open reading frame and leading to a premature stop codon which produces a truncated protein without the entire kinase domain. The correct integration of the targeting sequence within the Pifraire1 gene was confirmed by Polymerase Chain Reaction (PCR) and Southern blot hybridization analysis in the 129/SvEvBrd embryonic cells that were transformed (Fig. S3B). Following Bgl II or Acc65 I restriction digestion of the genomic DNA, Southern blot was performed using probes targeting 5’ or 3’ regions outside of the targeting vector, respectively. The predicted band sizes of 15.9 kb after Bgl II digestion with the 5’ probe and 13.5 kb after Acc65 I digestion with the 3’ probe confirms the homologous recombination of the external cassette. The transformed PFTK1 allele was easily detectable in the transformed mice through the agouti coat color of the chimeras. The produced chimeric mice were viable and fertile; after being crossed with wild type mice they also produce the PFTK1 heterozygote deficient mice that were viable and fertile. PFTK1 F1 generation, homozygote (null) mice, which were obtained by breeding heterozygous parents were also viable and fertile. To establish PFTK1 knockout line on a pure C57BL6 background, we backcrossed mixed (129/SvEvBrd x C57BL/6) heterozygote mice from the PFTK1 deficient line with C57BL6 for 10 generations, which continue to be viable. PCR performed on tail genomic DNA samples obtained from PFTK1 wildtype, heterozygotes and knockout PFTK1 mice and the corresponding primer pairs (fwd. 5’-TGACCCTTGTCTTTGAATACGTG-3’ and rev. 5’- GGAAGCCTAAATTGTAAGGATCAGG-3’ generating a 418 bp wild-type fragment and fwd. 5’– GCAGCGCATCGCCTTCTATC-3’ and rev. 5’-GGAAGCCTAAATTGTAAGGATCAGG-3’ generating a 341 bp fragment) are presented in Fig. S3C.

PCR mix was run with GoTaq Green Master Mix 2x (Promega, M712B). The PCR amplification process was as follows: initial amplification designed as touchdown PCR including: 6 cycles of denaturation 98 °C for 17 sec, annealing 69 °C (touchdown, -1°C per cycle) for 15 sec, amplification 72°C for 15 sec, and further amplification of the target genes was as follows: 35 cycles of 98 °C for 17 sec, 60 °C for 15 sec, and 72 °C for 15 sec. PCR products were resolved on a 1.5 % agarose gel-Ultra Pure Agarose (Invitrogen, 15510-027) and visualized by ethidium bromide (Ethyl hexadecyl dimethyl bromide C20H44BrN) (Sigma Ultra, C5335-1000).

### Real Time PCR (Quantitative RT- PCR)

All steps were performed in RNAase-free environment. To ensure RNA-free surface, RNase away (MBP Molecular Bioproducts, 7003) was applied. Total RNA was extracted from mouse embryo whole brains/cortices at embryonic day E 13.5-14.5 using Trizol Reagent (Ambion, 15596026) based on Invitrogen protocols. 25 ng of extracted RNA was used per reaction. To determine disruption of PFTK1 transcription the SuperScript III Platinum SYBR Green One-Step RT-PCR kit (Invitrogen, 204174) was used. Negative controls, no RNA and no RT were run in parallel with both the GAPDH housekeeping gene and PFTK1. Each sample was run in quadruplets and n=3. Results were normalized against GAPDH. Primers used for amplification of the target genes were as follows: (GAPDH-Fwd.: 5’-GGTGAAGGTCGGTGTGAACG-3’, GAPDH-Rev: 5’- C T C G C T C C T G G A A G AT G G T G - 3 ′) to generate a 233 bp product and (PFTK1 - Fwd: 5 ′ - CAGCGATCTGCCTCCACGGC-3’) designed within exon13. (PFTK1-Rev: 5’-GGCCCTCATGCTCTCTCCAGC-3’) designed within exon 14, to generate a 341 bp product. The primers would also be able to produce a short heptamer within mutated region (exon 8); thus, it could not interfere with mRNA levels for PFTK1-KO. The program for RT-PCR amplification was as follows: 48 °C for 30 min, 95 °C for 10 min, and 40 cycles of 95 °C for 15 sec, 60 °C for 30 sec and 72°C for 30 sec. The RT-PCR products were resolved on a 1.5 % agarose-ethidium bromide gel as well.

### Cell Culture

Human Embryonic Kidney cells (HEK-293T) were cultured in Dulbecco’s Modified Eagle Medium (DMEM) (High Glucose 4500 mg/L with 4 mM L-Glutamine without Sodium Pyruvate 0.1 µm sterile filtered) (ThermoScientific, SH30022-01) supplemented with 10 % Fetal Bovine Serum (FBS) (Hyclone, SH30397.03) and 1 % Antibiotic/ Antimicotic Solution100x (10,000 units/ml of penicillin, 10,000 µg/ml of streptomycin, and 25 µg/ml of Amphotericin B, 0.2 µm filtered) (ThermoScientific, SV30079.01) at 37°C in a 5% CO2 atmosphere.

### Primary Cortical Neuron Cultures

Cortical neuron cultures were prepared either from wild-type CD1 mouse embryos purchased from Charles River or from heterozygous crosses between PFTK1 mice, at gestational day /embryonic day E 13.5-14.5. Embryos were considered E 0.5 d, at the time vaginal plug was detected. Embryos were removed from the uterine horns and transferred to Hanks’ Balanced Salt Solution (HBSS) (modified without Calcium and Magnesium) (Hyclone Gibco, SH3003102). Brains were excised by cutting the skull open at the midline, from base of the head towards the front. Two lateral incisions were made under the cerebral cortices and skull was peeled off, meninges were removed and individual cortices were transferred to HBSS containing 8 µl Trypsin (25g/L in 0.9 % NaCl) (Sigma, T4549), incubated for 20 min at 37 °C with agitation. 10.7 µl of Trypsin inhibitor in neurobasal medium was added to stop trypsinization, and 13.34 µl DNase1 (10 mg / ml) (Boehringer Mannheim Roche Diagnostics, 10137100) to degrade extracellular DNA medium were added to the solution. Cells were pelleted at 1000 x, for 5 min at 4 °C and 10.7 µl of trypsin inhibitor and 13.34 µl of DNase were added. Cells were triturated; viable cells were counted using sterile filtered trypan blue solution (0.4 %) (Sigma, T8154). (Number of cells per ml was calculated by: cell count in the central square x dilution factor x 104). Cells were diluted to desired concentration for culture, in Complete Neurobasal Medium containing: Neurobasal medium (Gibco LifeTechnologies, 21103-049), B27 supplement (Invitrogen, 17504-044), N2 supplement (Invitrogen, 17502-048), L-Glutamine 200 mM (Gibco, 25030-081) and Penicillin-Streptomycin. Primary cortical neuron cells were plated into 24 or 6 well plates. Plates were coated with Poly-D-Lysine-Hydrobromide (1 mg/ml, sterile) (Sigma, P0899) prior to use. Poly-D-Lysine incubation was performed for at least 30 min and then plates were washed. For immunostaining, sterilized coverslips were placed into 24 well culture dishes and then coated with poly-D-lysine. Cultures were incubated at 37 °C in a 5% CO2 atmosphere and fixed or lysed at varied times.

### Expression Vectors

Plasmids for transfection: GFP subcloned into pCig2 vector (generously gifted by Carola Sherman), FLAG-Tagged: WT- PFTK1 and D228N- PFTK1 cDNA (mutation of conserved aspartic acid residue to asparagine, by a point mutation) (1284 kb) subcloned into pCMV vector (generously gifted by Dr. Paul Albert and Maribeth Lazzaro) (Lazzaro MA, 1997); GST-Tagged: WT-Rac1, G12V-Rac1, S17N-Rac1 subcloned in pEBG vector; GST-tagged: WT-Cdc42, G12V-Cdc42, S17N-Cdc42 subcloned in pEBG vector and MYC-tagged: WT-RhoA, CA-RhoA Q63L, DN-RhoA L19N in pRK5 vector. Rho GTPases (generously gifted by Dr. Gary Bokoch and Dr. Margaret Chou, University of Pennsylvania). Constructs were sequenced at Ottawa Genomic Center, using following primers: (1) For Rac1 and Cdc42 constructs in the pEBG backbone: (pGEX 5’ Sequencing Primer: 5’- [G G G C T G G C A A G C C A C G T T T G G T G] - 3 ’ and p GEX 3’ Sequencing Primer 5’ - [CCGGGAGCTGCATGTGTCAGAGG]-3’) (2) To sequence PFTK1, CMV virus backbone: (pCMV 5’ Sequencing Primer: 5’-CGCAAATGGGCGGTAGGCGTG-3’ PFTK1 GC rich region at 3’ end: 5’- GGGGGAGTTGAAGCTGGCAG-3’). Sequencing was performed 100 bp upstream of start codon and 100 bp downstream of stop codon. Plasmids were transformed with Top10 competent cells. Transformed cells were cultured on Ampicillin resistant agar plates (LB Agar Granulated, Molecular Genetics BP-1423-500). A single colony was picked and grown in Luria-Bertani (LB) Broth (Miller Amresco, J-106-500G). DNA was purified using the Pure Yield Plasmid Midi Prep System (Promega, A2495).

### Transient Transfection with Lipofectamine

HEK 293T Cells were cultured to 70% confluency, then co-transfected with Flag tagged (WT- PFTK1 or D228N- PFTK1) and GST tagged [WT-Rac1, G12V-Rac1 (constitutively active), S17N-Rac1 (dominant negative inactive)] or GST tagged [WT-Cdc42, G12V-Cdc42 (constitutively active), S17N-Cdc-42 (dominant negative inactive)] or GST tagged [WT-RhoA, Q63L-RhoA (constitutively active), L19N-RhoA (dominant negative inactive)], as well as, the reporter vector (GST::PEGB, MYC::PRK5, Flag::CMV using lipofectamine 2000 reagent (1mg/ml) (Invitrogen, 11668-019). Opti-MEM (Minimal Essential Medium), 1x Reduced Serum (Gibco, 31985-070) was used to dilute the plasmid and lipofectamine according to the manufacturer’s instructions. Diluted plasmid and lipofectamine were mixed and incubated for 20 minutes at room temperature. Cell culture medium was changed to media without FBS at the time of transfection. The plasmid-lipofectamine complex was added to culture media in droplets. 24 hours post transfection, cells were triturated off culture plates by scraping, cells were collected into ice cold PBS and spun down at 1.4 x g for 1 min for pull down assay.

### Viral Infection

Gene delivery via Adenoviral (AV) infection was performed at the time of plating. Adenovirus expressing WT- PFTK1 (Ad ET-CMV-3x Flag-WT- PFTK1, Pfu: 1.00 × 1012 CsCl Pure, Plaque pure), D228- PFTK1 (Ad ET- CMV-3x Flag-D228N- PFTK1, Pfu: 9.20 × 1011 CsCl Pure, Plaque pure), and GFP (Ad ET-CMV-EGFP, Pfu: 4.89 × 1011 CsCl Pure, 3x Plaque pure), was mixed with neuronal cortical culture cells at 50 -100 MOI (multiplicity of infection). Cells were fixed or lysed, at different time points, post-culture, for further analysis, depending on the nature of the experiment.

### Neuronal Death Assay

Cortical culture cells from mouse embryos at E13.5-14.5 were seeded at (1 × 105) in 96 well plates and infected/ co-infected with AV viruses (GFP alone or GFP + WT- PFTK1, D228N- PFTK1) at MOI=100 at the time of plating, for survival assay. Cells were washed with 1X PBS and then fixed with 4% PFA 12, 16 and 24 hours post-infection and then stained with Hoechst 33258 (1:10000) (Sigma-Aldrich B2883) in 1x PBS and incubated for the minimum of 30 min for nuclei staining. Intact nuclei were counted against condensed and fragmented nuclei to determine live neurons versus dead ones. To determine survival the number of live GFP positive neurons were assessed in each well, each treatment was performed in triplets, and the average for each treatment was calculated. Survival was assessed for WT- PFTK1 or D228N- PFTK1 as the percent of live GFP positive neurons in comparison to GFP. Survival percentage represents the ratio of GFP expressing neurons with morphologically intact nuclei at (12, 16, and 24 h) in WT- PFTK1 or KO- PFTK1 to the total number of GFP expressing neurons. The values were normalized to GFP alone as control. Data are presented as (Mean± SEM) of at least three independent experiments.

### Total Protein Extraction

To compare protein expression levels, whole brain lysates were prepared from mouse brains at E13.5-14-5 or from mouse pups at P21. Basically, mouse brains were dissected and washed with ice-cold PBS. Brains were homogenized in lysis buffer: [50mM Tris-HCl (pH=7.5), 150mM NaCl,1 mM MgCl2, 1mM EDTA, 1% triton X-100 (Boehringer Mannheim, 789704), 1mM L-Dithiothreitol (DTT, C4H10O2S2) (Sigma Aldrich, D0632-5G), protease inhibitor (Halt Protease inhibitor Cocktail EDTA-free 100x (Roche, 87785)] for 20 min, at 4 °C, with agitation. Lysates were clarified by centrifugation at 19500 x g, 4 °C, for 20 min. Supernatant was collected (approximately 2mg of protein was collected per brain), mixed with 0.1% w/v bromophenol blue/ DTT sample buffer and heated at 100°C, for 10 min.

### Immunoprecipitation (IP)

All steps were performed on ice. Cell lysate was prepared as described, previously. Lysate was prepared using lysis buffer 50 mM Tris-HCl (pH=7.5), 150 mM NaCl, 1 mM MgCl2, 1mM EDTA, 1% Triton X-100 (Boehringer Mannheim, 789704), 1mM DTT (L-Dithiothreitol, C4H10O2S2) (Sigma Aldrich, D0632-5G), protease inhibitor, for 20 min, at 4°C with agitation. Lysates were clarified by centrifugation at 19500 x g, 4°C, for 20 min. Supernatant was collected and incubated with 20 µl of washed bead per reaction. After immunoprecipitation, beads were pelleted at 1000 x g, washed five times with wash buffer (Tris-HCl (pH⁈7.3-7.5), NaCl, EDTA, Triton 10 %, DTT), then mixed with 0.1 % w/v bromophenol blue / DTT sample buffer, and heated at 100°C, for 10 min to release the protein complex. Proteins were resolved on 12 % SDS-polyacrylamide gels.

For immunoprecipitation of ectopically expressed GST tagged proteins, GST beads - (Glutathione Sepharose 4b beads, particle size 45 µm - 165 µm, GE Healthcare 17-0756-05) were incubated with the lysate overnight, at 4 °C, on a rotor. For immunoprecipitation of endogenous protein, Trueblot Anti Rabbit IgIP beads - (eBioScience, 00-8800-25) or Anti mouse IgIP beads - (eBioScience, 00-8811-25) were used, depending on the origin of the bait antibody. Lysate was collected and precleared by incubating 20 µl of the washed bead that was to be used for IP, for 30 min, at 4 °C, on a rotor. Supernatant was cleared by centrifugation at 1000 x g, 1 min, at 4°C. 4 - 8 µg of antibody per reaction, normal mouse IgG (SantaCruz, sc-2025) or normal rabbit IgG (SantaCruz, sc-2027) and Pftaire-1 (H-140) (1:500) rabbit polyclonal IgG (SantaCruz, sc-50475), RhoA (1:500) mouse monoclonal IgG (SantaCruz, sc-418), Rac1 (C-11) (1:500) rabbit polyclonal IgG (SantaCruz, sc-95), Cdc42 (1:500) rabbit polyclonal IgG (SantaCruz, sc-87) was incubated with the cleared supernatant, overnight, at 4°C, on a rotor. PFTK1 was used to IP GTPase and vice versa. 20 µl of the washed beads were incubated with the antibody-lysate mix for 3 hours, at 4 °C, on a rotor to precipitate the antibody.

### SDS-PAGE and Western Blot

Protein extracts were resolved on a 10 or 12 % SDS - polyacrylamide gel, along with Molecular Marker, Blueye Prestained Protein Ladder (GeneDirex, P14007-0500). Transfers were done on 0.45 µm PolyVinylidene DiFluoride (PVDF) Transfer membrane (Immobilon-P, IPVH00010). Membrane was blocked in 10 % non-fat dry milk in 1 x PBST [Phosphate Buffered Saline (pH=8.0) and 0.1 % Tween] at room temperature, for 1 hour. Proteins were detected using antibodies. Primary antibodies were diluted in [3 % Bovine Serum Albumin (BSA) fraction V (Fisher, BP1600-100) in PBST and 0.1 % Sodium Azide NaN3 (Sigma Ultra, s-8032)] as follows: PFTK1 (H-140) (1:500) rabbit polyclonal IgG, (SantaCruz, sc-50475), Pftaire2 (CDK15) (1:500) rabbit polyclonal (Life Span Biosciences, LS-C119219/34326), RhoA (1:500) mouse monoclonal IgG (SantaCruz, sc-418), Rac1 (C-11) (1:500) rabbit polyclonal IgG (SantaCruz, sc-95), Cdc42 (1:500) rabbit polyclonal IgG (SantaCruz, sc-87), Cdk5 (C-8) (1:500) rabbit polyclonal IgG (SantaCruz, sc-173), Pctaire1 (CDK16) (1:500) rabbit polyclonal, (Life Span Biosciences, LS-C112292/34325), Anti-Flag (1:500) rabbit monoclonal IgG (Sigma, F7425), Anti-GST (1:500) mouse monoclonal IgG (SantaCruz, sc-138) and incubated with the membranes overnight, at 4 °C, with agitation. Membranes were washed and then incubated with secondary antibodies diluted in 5 % non-fat milk, for 1 hour, at room temperature [goat anti-mouse 170-6516 or goat anti-rabbit 170-6515 IgG (H+L) HRP conjugate (Bio-Rad)] were all used at 1:2000 in 3 % BSA in 0.1 % Tween 20 (PBS/T). Immunoreactivity was detected using Immobilin Western Chemiluminescence HRP substrate (Millipore, WBKLS0500) or Pierce ECL Western Blotting Substrate (Thermo Scientific, 32106) and visualization HyBlot CL Autoradiography film, Denville Scientific. Protein levels were normalized against β-actin signals, and results were reported in reference to control values (untreated control for each individual experiment).

### PFTK1 Kinase Assay

Kinase assay was performed using immunoprecipitated WT- PFTK1 protein or IgG as a control. Briefly, Neuronal cortical cells prepared from WT- PFTK1 mouse embryos E13.5-14.5 were cultured in Neurobasal media (Gibco, 21103-049) containing B27 (Invitrogen, 17504-044) and N2 (Invitrogen, 17502-048), in 12 well plates, at the ratio of 1 embryo per well, for 18-20 hour, at 37°C, 5 % CO2. After washing with ice-cold 1x PBS, lysates were collected directly from plates by scraping into ice-cold immunoprecipitation (IP) buffer (Tris-HCL 25mM (pH 7.4), Na4P2O7 10 mM, Na3VO4 25 mM, NaF 50 mM, EDTA 1 mM, Digitonin 2 mg/ml) and sonicated twice, for 3 cycles, with a 20 minute incubation in between the 2 set of cycles, and then supernatant was cleared by centrifugation at 14000x g, 4°C, for 10 min. 1 mg of protein(adjusted to the volume of 300 µl was incubated with 4 µg antibody either Normal rabbit IgG (200 µg/0.5 ml) or PFTK1 (200 µg/ml) antibodies, overnight, at 4°C, on the shaker. IP-ed, WT- PFTK1 and IgG were mixed with 40 µl of pre-equilibrated (performed washes with IP wash buffer) Trueblot anti-rabbit IgIP beads per reaction and incubated for 2 hours, at 4°C, on the shaker. Beads were then washed and pelleted at 2000rpm, for 1 min, at 4°C, with IP wash buffer and kinase assay buffer [HEPES 50 mM (pH 7.5) and 1mM DTT], respectively. Next, kinase reaction buffer (HEPES 50 mM (pH 7.5), 1 mM DTT, 10 mM MgCl, and 20 µM ATP)) was added to a final volume of 25 µl. 10 µg of RhoA substrate His-Recombinant human RhoA (Prospec, pro-057), was added to reactions with or without (5 µl, 100 x) GDP. Reactions were incubated overnight, at 30°C. Samples were then centrifuged at 2000 rpm, for 1 min, at 4°C, then supernatant was collected and subjected to either western blot for rabbit polyclonal IgG phosphoserine (Invitrogen, 61-8100) signal; or G-LISA assay to assess RhoA activity as described below.

### Assessment of Rho GTPase Activity

Activation of RhoA, as a function of PFTK1 kinase activity was determined using commercially available “RhoA Activation Assay Kit (CellBioLabs, STA-403-A)” according to the manufacturer’s instructions. Primary Cortical Culture Cells were prepared from Het x Het crosses at embryonic day 13.5-14.5, at the density of 1 embryo per well, in 1.5 ml, cultured in complete Neurobasal media supplemented with B27 and N2, in 12 well plates, and incubated at 37 °C and 5 % CO2. 18-20 hours post-plating cells were washed with ice-cold PBS and GTPase assay was performed according to the manufacturer’s kit. To decrease rapid hydrolysis of active GTP bound to inactive GDP bound protein all following procedures were performed on ice. Briefly, cells were incubated on ice, for 20 minutes, with 1x Assay / lysis buffer provided with the kit (Part No. 240102) (containing 125 mM HEPES, pH 7.5, 750 mM NaCl, 5% NP-40, 50 mM MgCl2, 5 mM EDTA, 10% Glycerol) and then collected by scraping. Due to very low expression of RhoA same-genotype extracts were pooled to achieve large amounts of Protein. Total protein was quantified by Bradford protein assay to achieve at least 800ug of protein for each sample/assay. Cell lysates were clarified by centrifugation at 14,000 g x, at 4 °C, for 10 min. (at least 800 µg protein per reaction). 100x GTPγS (Part No. 240103) and GDP (Part No. 240104) were added to the cells for positive and negative control, respectively. Control samples were incubated for 30 minutes, at 30 °C, with agitation, and the reaction was terminated by adding 1M MgCl2 to the samples. Volume of sample per reaction was adjusted to be identical in all reactions and 40 µl of bead added per reaction. Active proteins were immunoprecipitated with respective rhotekin RBD agarose beads for RhoA. Beads were pelleted by centrifugation at 14000 x g, for 10 seconds, and washed three times with 1x Assay buffer supplied in the kit. Afterwards, beads were resuspended in SDS-PAGE sample buffer and boiled for 5 minutes for detection by western blotting. Separated proteins were visualized using RhoA, Mouse Monoclonal antibody (Part No. 240302) provided with the kit.

To quantify Rho GTPase activity after the PFTK1 Kinase assay, in a more accurate/sensitive manner, G-LISA RhoA activation assay biochem kit (Absorbance based) (Cytoskeleton, BK124) was used according to the manufacturer’s protocol. Samples from the kinase reactions, (IgG-IP: RhoA, IgG-IP: RhoA+GDP, PFTK1-IP: RhoA and PFTK1-IP: RhoA+GDP) were diluted and pre-equalized with ice cold lysis buffer /binding buffer (Part #GL37). To pre-equalize samples, protein concentration was determined at 600nm, using Precision RedTM Advanced Protein Assay Reagent (Part # GL50) as suggested by the manufacturer. Same buffer mix (no protein) and RhoA (provided by manufacturer), were respectively, used as blank and positive control for the assay. Next, samples were transferred to 96 well Rho-GTP binding plates (Part #GL25), equipped with pre-washed GLISA assay strips, incubated at 4°C, on an orbital microplate shaker at 400 rpm, for 30min. Samples were then washed with wash buffer (Part # GL38) and incubated with Antigen Presenting Buffer (Part#GL45) for 2 mins, washed again with wash buffer, then incubated at room temperature for 45 min, on orbital microplate shaker at 400rpm, with anti-RhoA primary antibody (Part # GL01) (1:250) in Antigen dilution buffer (Part #GL40) and washed subsequently. Samples were next incubated with (1:62.5) secondary horseradish peroxidase (HRP) conjugate antibody (Part #GL02) at room temperature, for 45 min, on an orbital microplate shaker at 400 rpm and washed afterwards. Then samples were incubated with HRP detection reagents A (Part # GL43) and B (Part # GL44) (1:1 ratio) for 15 min, at 37 °C, and reaction was stopped by adding HRP Stop Buffer (Part #GL80), and absorbance measured at 490 nm using a microplate spectrophotometer. To measure RhoA activity, values were determined as follows, 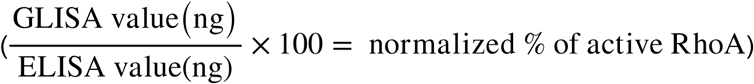. The detected signal was normalized against total RhoA using G-LISA and E-LISA (BK#150) kits side by side. To determine if any significant difference existed in the activity of RhoA among samples, two-way ANOVA was performed in the grouped data followed by Bonferroni’s post-hoc test. Results were presented as average of at least three independent experiments.

### Brain Sectioning

Brains were collected from the Wt- PFTK1 and KO- PFTK1 mice at 8 weeks of age. Animals were humanely anesthetized with euthasol and transcardially perfused with 0.9 % saline, followed by fixative solution 4 % Paraformaldehyde (PFA) (EMD, PX0055-3) in 1x PBS at pH=7.4. Brains were then removed from the skull and fixed further with 4 % PFA, at 4 °C, for 24 hrs. Fixed brain tissue was cryopreserved in 20 % sucrose in 0.1 M phosphate buffer and stored at 4°C, over a 3-day course, and the solution was changed 3 times per day. Brains were then snap-frozen with Nitrogen balanced with CO2 and then embedded with O.C.T. (Tissue - Tek /O.C.T compound, 4583) for sectioning. 14 µm coronal sections were collected from the brains; starting at the frontal cortex at Bregma: ∼ 2.710 mm up to the 4th ventricle at Bregma: ∼ - 4.20 mm. For collecting, 3 sections were skipped in between each selection. Sections were directly mounted onto SuperFrost Plus microscopic slides 25 × 75 × 1.0 mm (Fisher brand, 12-550-15) and subjected to2 % cresyl violet staining as described below.

### Cresyl Violet Staining

Sections were thawed on slide warmer for 15-20 min, at 35 °C, and then placed in distilled water for 5 min. Sections were then stained with 2 % filtered cresyl violet (pH=3.5) cresyl violet acetate (C18H15N3O3, Sigma C5042-10G), for 10-30 minutes depending on the stain concentration. Sections were then dehydrated in graded ethanol solutions from 50 % to 100 %, followed by defatting with xylene. Sections were mounted with coverslips using toluene.

### Immunostaining

Cells plated on glass coverslips were fixed in 4 % PFA in 1 x PBS, for 15 min. Fixed cells were permeabilized and blocked simultaneously in 0.1 % Triton X-100, and 5 % BSA in 1x PBS, respectively, for 20 min, at room temperature. Coverslips were incubated with primary antibodies diluted in 5 % BSA in 1x PBS: Tau1 clone 46 (1:500) mouse monoclonal IgG (Sigma-Aldrich, T-9450), GFP (H-140) (1:500) mouse monoclonal IgG (Abcam, ab1218), PFTK1 (H-140) (1:500) rabbit polyclonal IgG (SantaCruz, sc-50475), MAP2 (H-300) (1:500) rabbit polyclonal IgG (SantaCruz, sc-20172) for 1hr, at room temperature. Coverslips were washed; incubated with FITC-secondary antibodies: Alexafluor 488 (1:1000) Goat-Anti-mouse (Molecular Probes, A11001), Streptavidin Alexafluor 488 (1:1000) Goat-Anti-mouse (Molecular Probes, S11223), Streptavidin Alexafluor 546 (1:1000) Goat-Anti-mouse (Molecular Probes, S11225), Alexafluor 594 (1:1000) Goat-Anti-mouse (Molecular Probes, A11005), Alexafluor 594 (1:1000) Goat-Anti-rabbit (Molecular Probes, A11037), Alexafluor 594 (1:1000) Donkey-Anti-Goat (Molecular Probes, A11058) were diluted in 5% BSA in 1x PBS, and then incubated with coverslips for 30 min, at room temperature. To detect intact nuclei, cells were stained with Hoechst 33258 (1:10000) (Sigma-Aldrich B2883) in 1x PBS, incubated for the minimum of 30 min, at room temperature. Slides were immersed in distilled water to remove excess salt and then mounted on slides applying approximately 25 µl of Vectashield Mounting Medium (Vector laboratories, H-1200).

### Microscopy and Imaging

Images were taken using upright Zeiss Axioskop 2 Mot or invert Zeiss AxioObserver-Z1 fluorescent microscope with Northern Eclipse Software or Zeiss and Axiovision Rel. 4.8 software, captured in XY mosaic format, in rectangle mode. Images were taken at 20x, 32x, and 40 x magnification lenses. To determine the total diameter of the field of view for each objective, Ocular Field of View (F.O.V.) number (32) was divided by objective magnification and following measurements were obtained: for 20x (1500 µm), for 32x (719 µm), and for 40x (575 µm). Images were processed for length measurements using Axiovision software Rel. 4.8, spline mode to trace axon length or Stereo Investigator microscopy imaging system (version 6; MicroBrightField, Williston, VT) software using contour mapping mode and tracing the length of axons. The distance from the cell body of the GFP expressing neuron or Tau1 to the distal region of the longest neurite was measured as axon/lengthiest neurite.

### Quantification of Signal Density on Images

Immunoblots were further analyzed by Image J software to compare the density/intensity of the bands of an agar gel or a western blot membrane. Following imaging of the blots, representative bands were selected, and their intensity was quantified. Then quantified measurements were transferred to excel and results were normalized against relative controls (β-actin for western blot and S12 for RT-PCR).

