## Supplementary material for "PFTK1 kinase regulates axogenesis during development via RhoA activation": Suppl Figures

| Stages | Eip63E$^{81}$ Axons | Eip63E$^{81}$ neurons | Df(3L)E1 Axons | Df(3L)E1 neurons |
|---|---|---|---|---|
| 8-9 | 10.00 | ND | ND | ND |
| 10-11 | 35.84 | 36.10 | 7.14 | 28.60 |
| 12-13 | 50.00 | 52.86 | 88.00 | 85.70 |
| 14-15 | 85.71 | 44.50 | 96.67 | 100.00 |

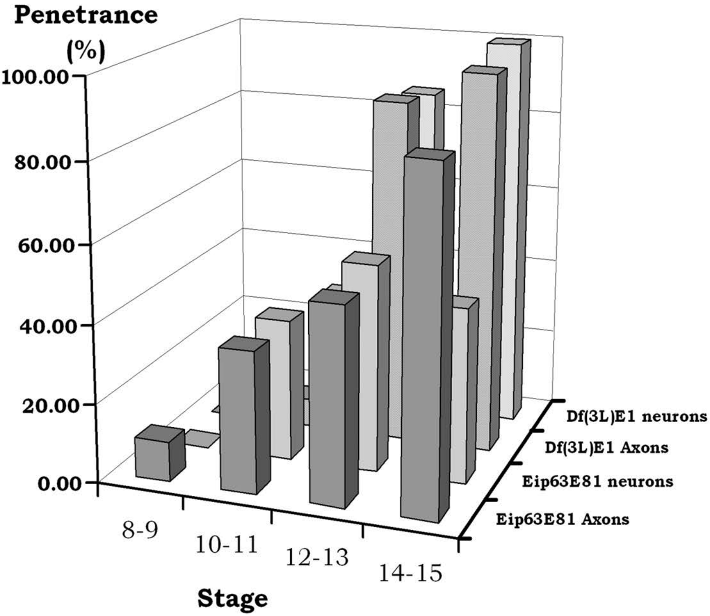


**Figure S1. Eip63E deficiency leads to concerted defects in axons and neurons of the Drosophila VNC.**
The combined penetrance of markers relevant to the same VNC structures are presented as one. Axon-associated penetrance reflects those calculated using HRP, BP-102 and Fasciclin-II antibodies. Neuron-related values include Neuroglian, Elav and Futsch-revealed penetrance. Penetrance was calculated as the % of defective KO embryos from the total KO embryos screened. Both deficient lines show the same nature of defects. Although it seems that defects associated to the Df(3L)E1 allele are more acute and appear later in development, Eip63E81 shows a robust axonal phenotype. ND: Not determined.

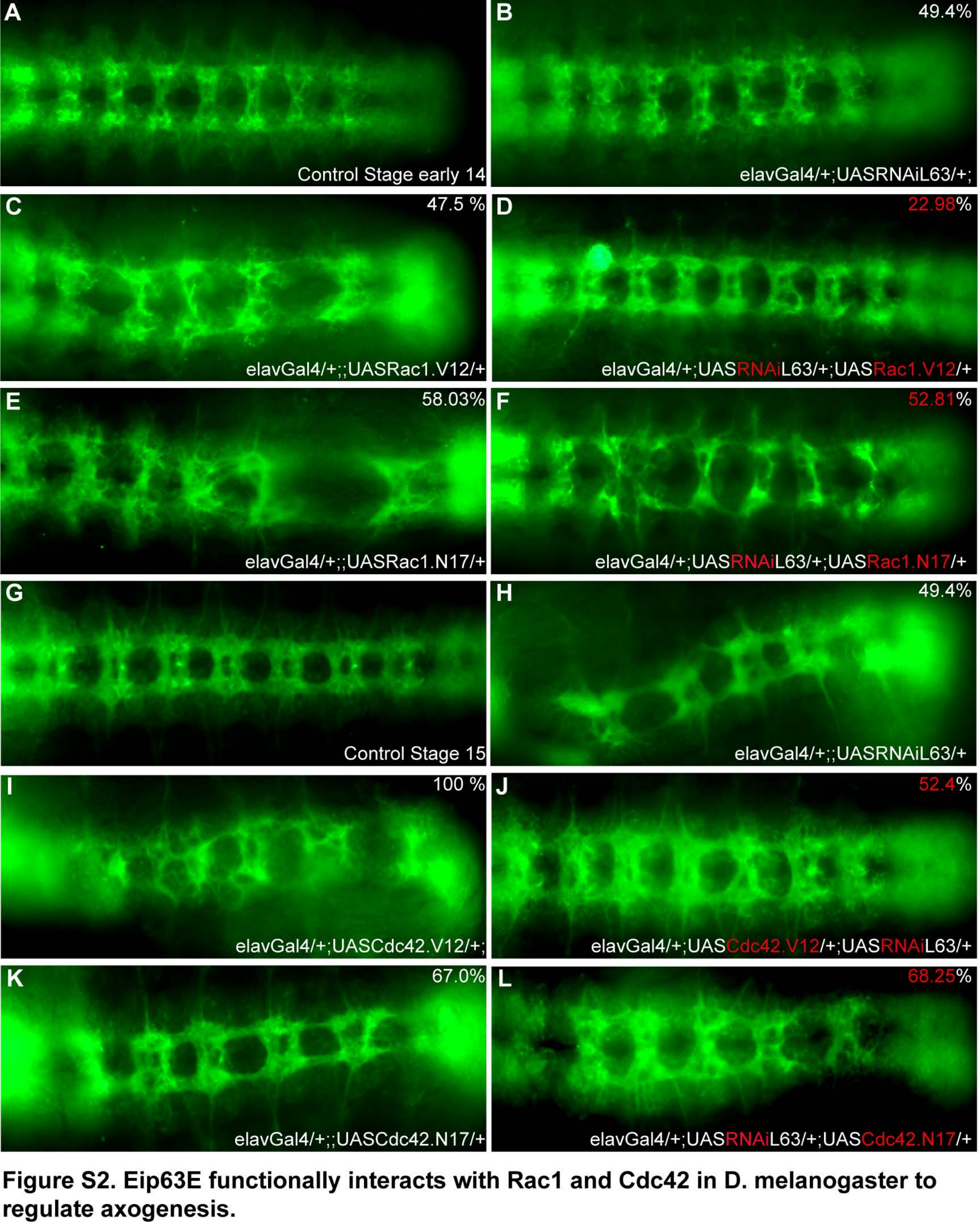


**Figure S2. Eip63E functionally interacts with Rac1 and Cdc42 in D. melanogaster to regulate axogenesis.**
Ventral views of stage 14-15 whole mount embryos stained for BP102 antigen are shown head to the left. Panels show representative images for each specified genotype. An UAS-elavGal4 approach was used to drive Eip63E downregulation and either constitutively active or dominant negative forms of Rac1 or Cdc42 overexpression in neurons of independent D mel. lines. Flies were then crossed to assess the nature of axon defects in embryos expressing both UAS constructs vs the single heterozygous controls. (A-F) Neuronal functional interaction between Eip63E and Rac1. Only the combination of Eip63E deficiency and constitutively active Rac1 (UAS Rac1 V12; D), but not DN-Rac1 (UAS Rac1 N17; F), leads to an intermediate phenotype. (G-L) Functional interaction between Eip63E and Cdc42. Ventral views of the indicated genotypes of crosses between Eip63E and Cdc42 constitutive active form (UAS Cdc42 V12) or the dominant negative Cdc42 (UAS Cdc42 N17) are presented.

A

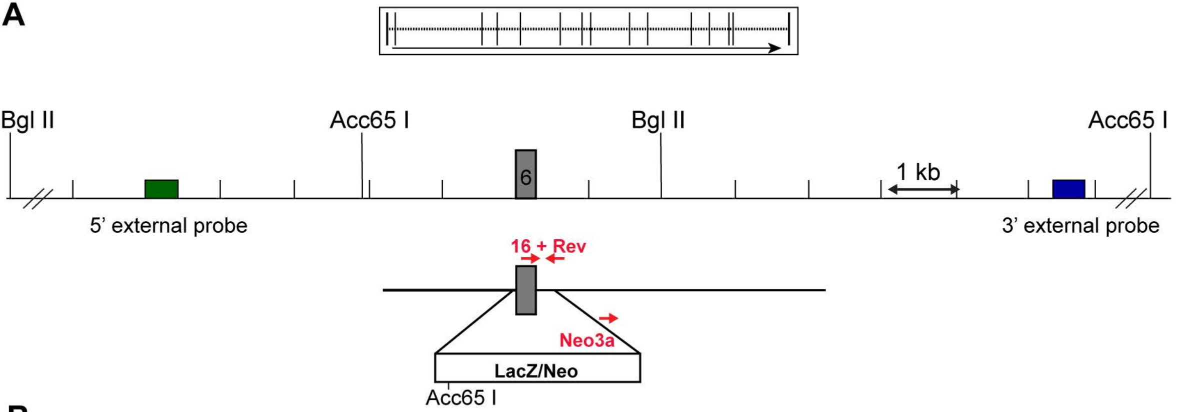


B

Southern Strategies

| Probe | 5' external | 3' external |
|---|---|---|
| Enzyme | Bgl II | Acc65 I |
| Wildtype | 10.9 kb | 11.2 kb |
| Targeted | 15.9 kb | 13.5 kb |

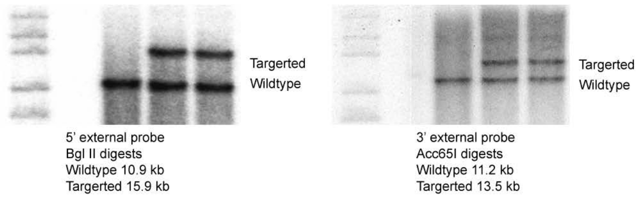


* Clone achieving germline transmission

C

PCRStrategies

| Stretegy | WT-specific | Mutant-specific |
|---|---|---|
| Primers | 16+Rev | Neo3a + Rev |
| Wildtype | 418 bp | --- |
| Targeted | --- | 341 bp |

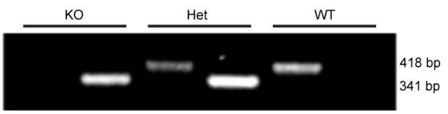


**Figure S3. Strategy for generation of Pftk1 (Cdk14) deficient murine line.**
The mouse was designed and developed by the Texas A&M Institute for Genomic Medicine (TIGM) (A) Strategy. Exon 6 (from nucleotide 18-Exon6 to nucleotide l-Exon7) of the Cdk.14 gene at chromosomes (4,803,391-5,380,19 7) was replaced for an IRES/bGeo/Poly A cassette by homologous recombination. (B) Southern Blot screening of clones. (C). PCR strategy for genotyping. Bottom panel shows representative gel image with samples from the three possible genotypes.

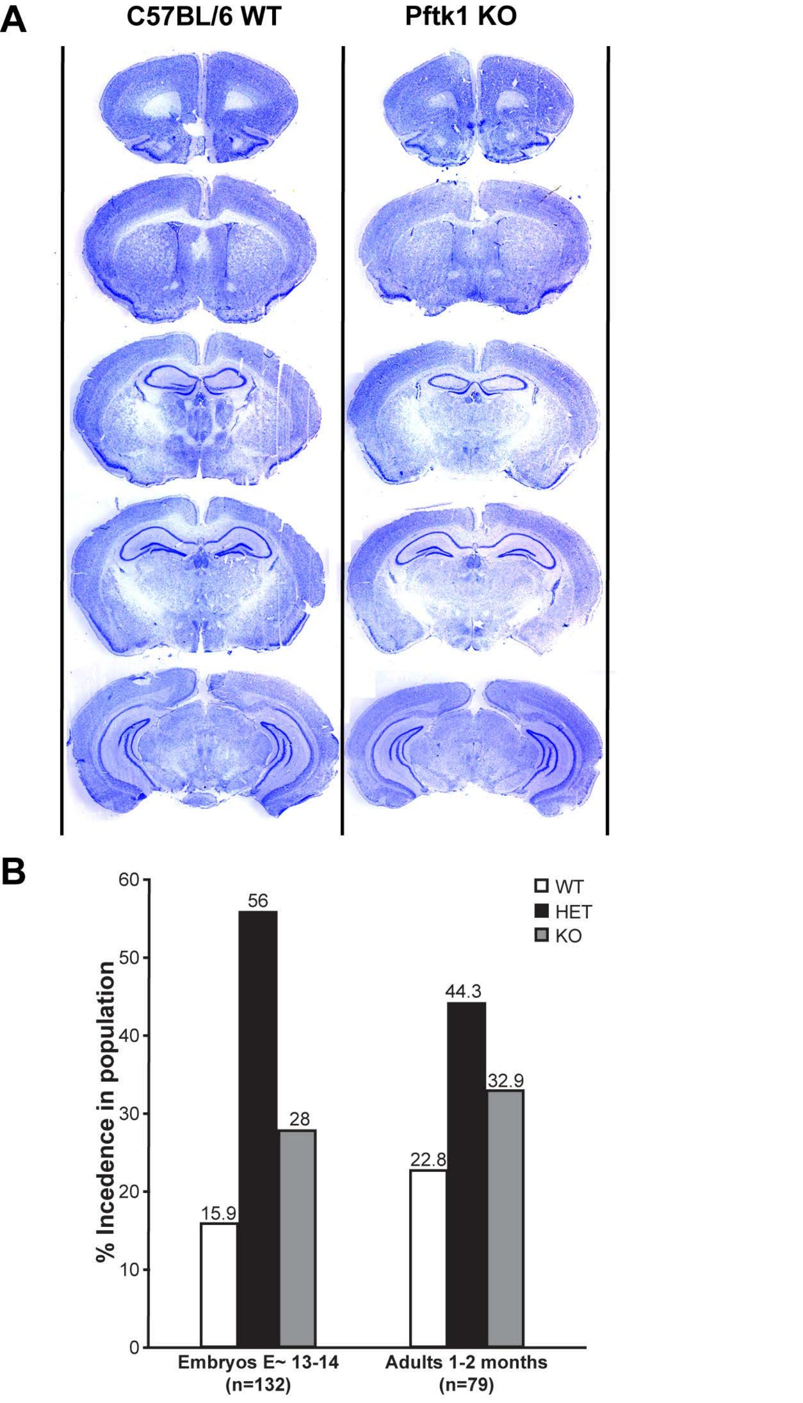


**Figure S4. Pftk1 deficiency does not appear to cause any gross brain defects or lethality.** (A) Brains from three-month-old mice (3rd generation backcrossed) were collected, perfused and fixed for sectioning. 14μm sections were collected from cryopreserved samples, starting from the cortex at bregma: ~ 2.710 mm up through the 4th ventricle at bregma: ~-4.20 mm. Sections were then stained with cresyl violet to examine the gross anatomy of the brain. No abnormality was detected for KO mice (n=3/genotype). Representative sections from bregma 1.98, 1.70, 0.74, 1.00, 1.64, and 2.75 mm are presented. (B). Embryos or mice (=> 6th generation backcrossed) from HET x HET crosses were genotyped and incidence of the three possible outcomes is presented as % of the total n. No gross deviation from the expected 25:50:25 Mendelian ratio was detected.

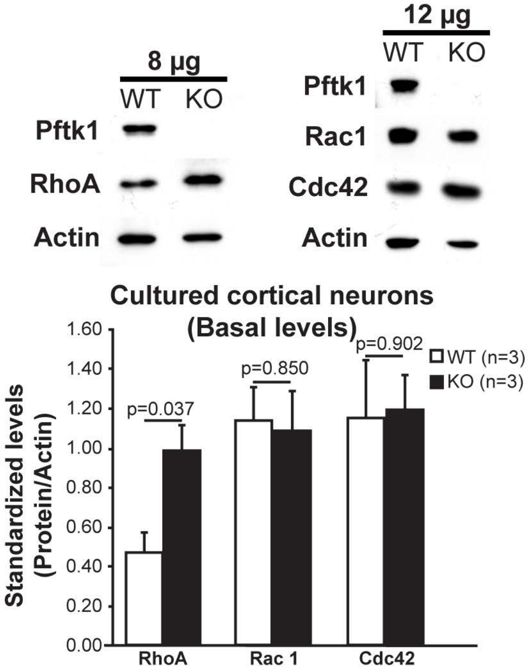


**Figure S5. Differential expression of Rho GTPases in brains of Wt vs KO E13-14 murine embryos.**

Only RhoA is misregulated in cortical neurons. Basal levels of RhoA, Rac1 and Cdc42 protein in WT and Pftk1-defficient cortical neurons. Primary cortical neurons from E13-14 embryos from HET x HET crosses were dissected and plated at 0.65-0.75x106 cells/ml. After 18-20h in vitro, proteins were extracted followed by western blot. Top: Representative images from western bot are shown. Bottom, levels were assessed by densitometry using Image J and Rho GTPases signals were standardized vs loading control (Actin). Each column represents the average from 3 embryos/genotype +SEM. Significant difference between WT and KO neurons was only found for RhoA levels. Top: Representative images from western blot are shown. Bottom, levels were assessed by densitometry using Image J and Rho GTPases signals were standardized vs loading control (Actin). Each column represents the average from n embryos/genotype +SEM. Significant difference between WT and KO neurons was only found for Cdc42 levels. t-test was used for statistical analysis whenever a p value is specified

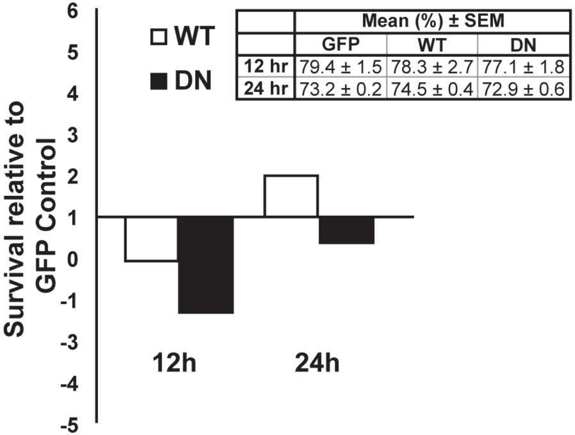


| | Mean (%) ± SEM | | |
|---|---|---|---|
| | GFP | WT | DN |
| 12 hr | 79.4 ± 1.5 | 78.3 ± 2.7 | 77.1 ± 1.8 |
| 24 hr | 73.2 ± 0.2 | 74.5 ± 0.4 | 72.9 ± 0.6 |

**Figure S6. Manipulation of Pftk1 levels has no impact on neuronal survival.**

Primary cortical neurons derived from CD1 embryos (E 14-15) were infected with 100 MOI of adenoviruses expressing GFP alone, GFP and WT-Pftk1 or GFP and D228N-Pftk1. After 12 or 24 hours in culture, cells were fixed and stained with Hoechst 33258 (nuclei visualization). nuclear integrity of GFP+ neurons was used as a criterion for vitality. Survival values represent the % of healthy nuclei of the total GFP+ cells. Table is showing average +SEM from three independent experiments. Graph shows the same data but presented as value relative to the GFP control.
